# Determinants of sugar-induced influx in the mammalian fructose transporter GLUT5

**DOI:** 10.1101/2022.06.17.495601

**Authors:** Sarah E. McComas, Tom Reichenbach, Darko Mitrovic, Claudia Alleva, Marta Bonaccorsi, Lucie Delemotte, David Drew

## Abstract

In mammals, glucose transporters (GLUT) control organism-wide blood glucose homeostasis. In human, this is accomplished by fourteen different GLUT isoforms, that transport glucose and other monosaccharides with varying substrate preferences and kinetics. Nevertheless, there is little difference between the sugar-coordinating residues in the GLUT proteins and even the malarial *plasmodium falciparum* transporter *Pf*HT1, which is uniquely able to transport a wide range of different sugars. *Pf*HT1 was captured in an intermediate “occluded” state, revealing how the extracellular gating helix TM7b has moved to break and occlude the sugar-binding site. Sequence difference and kinetics indicated that the TM7b gating helix dynamics and interactions likely evolved to enable substrate promiscuity in *Pf*HT1, rather than the sugar-binding site itself. It was unclear, however, if the TM7b structural transitions observed in *Pf*HT1 would be similar in the other GLUT proteins. Here, using enhanced sampling molecular dynamics simulations, we show that the fructose transporter GLUT5 spontaneously transitions through an occluded state that closely resembles *Pf*HT1. The coordination of fructose lowers the energetic barriers between the outward and inward-facing states, and the observed binding mode for fructose is consistent with biochemical analysis. Rather than a substrate binding site that achieves strict specificity by having a high-affinity for the substrate, we conclude GLUT proteins have allosterically coupled sugar binding with an extracellular gate that forms the high-affinity transition-state instead. This substrate-coupling pathway presumably enables the catalysis of fast sugar flux at physiological relevant blood-glucose concentrations.

## Introduction

Glucose (GLUT) transporters facilitate the rapid, passive flux of monosaccharides across cell membranes at physiologically relevant concentrations ranging from 0.5 to 50 mM ^1,2^. In human, most GLUT isoforms transport D-glucose, but with different kinetics, regulation and tissue distribution ^1,2^. For example, GLUT1 is a ubiquitously expressed transporter with saturation by D-glucose around 5 mM to maintain blood-glucose homeostasis, whereas the liver isoform GLUT2 is saturated at 50 mM, enabling a high-flux of glucose after feeding-induced insulin secretion ^1,3^. Others, like GLUT4, are localized to intracellular vesicles, but will traffic to the plasma membrane of adipose and skeletal muscle cells in response to insulin signalling ^4^. GLUT5 is the only member thought to be specific to fructose, and is required for its intestinal absorption ^5,6^. In this process, glucose is actively absorbed by sodium-coupled glucose transporters, while fructose is taken up passively by GLUT5 ^7^. GLUT5 must therefore efficiently transport fructose at high sugar concentrations (K_M_ = 10 mM), whilst still maintaining sugar specificity ^7^. It is poorly understood how GLUT proteins retain strict sugar-specificity and how sugars are able catalyse large conformational changes when they bind to GLUT proteins with weak (mM) affinities ^8^. As gate-keepers to metabolic re-programming ^9,10^, answers to these fundamental questions could have important physiological consequences for the treatment of diseases, such as cancer and diabetes.

GLUT transporters belong to the Major Facilitator Superfamily (MFS), whose topology is defined by two six-transmembrane (TM) bundles connected together by a large, cytosolic loop (Fig. 1A) ^11^. Within the MFS, GLUT proteins belong to a separate subfamily referred to as sugar porters, which are distinct from other well-known sugar transporters such as LacY ^8,12^. Sugar porters are subclassified based on a unique sequence motif ^13,14^, and crystal structures reveal that this motif corresponds to residues forming an intracellular salt-bridge network, linking the two bundles on the cytoplasmic side (Fig. 1A, Fig. S1) ^14,15^. The salt-bridges are formed between the ends of TM segments and an intrahelical bundle (ICH) of four to five helices. Crystal structures of GLUT1 ^16^, GLUT3 ^15^, and GLUT5 ^14^ and related homologues ^17-21^ have shown that the GLUT proteins cycle between five different conformational states: outward-open, outward-occluded, fully occluded, inward-occluded and inward-open (Fig. 1B). Whilst GLUT proteins are made up from two structurally-similar N-terminal (TM1-6) and C-terminal (7-12) bundles, structures have shown that glucose is not coordinated evenly, but almost entirely by residues located in the C-terminal bundle ^15^. In particular, residues in the half-helices TM7b and TM10b make up a large fraction of the sugar-binding site ^8^. The current working transport model is that the half-helices TM7b and TM10b undergo local conformational changes in response to sugar binding and control substrate accessibility to its binding site from the outside and inside, respectively (Fig. 1B)^11,14^. In brief, upon sugar binding from the outside, the inward movement of the extracellular gating helix TM7b is followed by the outward movement of TM10b, and the subsequent breakage of the cytoplasmic inter-bundle salt-bridge network, enabling the two bundles to move around the substrate ^8^. In the inward-facing state, TM10b moves fully away from TM4b, sugar exits, and the protein spontaneously resets itself to the outward-open state (Fig 1B). Resetting back through an empty-occluded state is rate-limiting and ∼100 fold slower than *via* a loaded-occluded intermediate ^2,22^.

**Fig 1.**
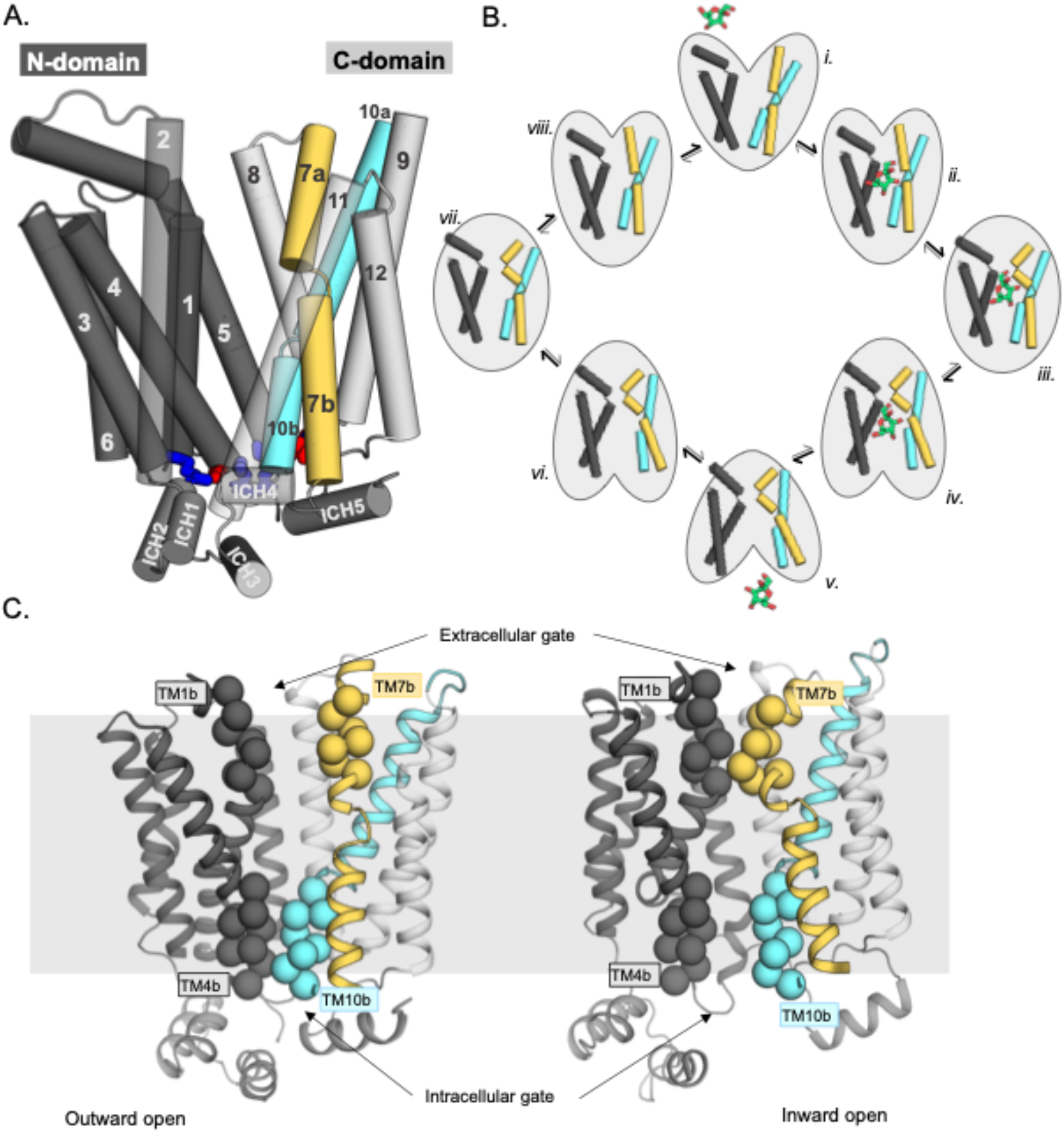
Schematic highlighting the GLUT structural features and their major conformations in the transport cycle. **A**. Structural overview of a sugar porter, GLUT5 (PDB:4ybq). The N-terminal bundle (left, dark grey, transmembrane helices 1-6) and C-terminal bundle (right, light grey, transmembrane helices 7-12) form the separate six-transmembrane bundles, which are connected by the large cytosolic loop comprising intracellular helices (ICH) 1-4. The salt bridge forming residues linking the two bundles are shown as sticks in blue and red, indicating the positive and negative charge of the side chains, respectively. The broken transmembrane helices TM7 (forming TM7a and TM7b) and TM10 (forming TM10a and TM10b) are colored yellow and cyan respectively. **B**. Schematic conformational cycle a sugar porter will undergo, based on currently available protein structures. Briefly, moving clockwise from top middle, the transporter will receive a sugar in the outward open state (i). The transporter then undergoes a partial occlusion of the extracellular gate (outward occluded, ii), followed by full occlusion (iii). Once both gates are fully shut, the inner gates begin to open (inward occluded, iv) where the salt bridge residues begin to lose contact. Finally, the salt bridges are fully broken apart in an inward open state (v), and the sugar can be released into the cell. The transporter will then go through the same motions in reversed order in the absence of sugar to reset to the outward open state (vi-viii). **C**. The extracellular gate is formed by TM1b and TM7b half-helices, and the intracellular gate is formed by TM4b and TM10b half-helices. Residues defining these gates are shown as spheres. In the outward open state (left), the extracellular gate is open, and intracellular gate is shut. In the inward open state (right), the opposite occurs. The grey slab behind the proteins indicates the rough location of the lipid bilayer membrane.

The intermediate, occluded conformation is arguably the most informative for understanding how sugar binding and transport are ultimately coupled ^8^. Due to its transient nature, this state is rarely seen. However, it was fortuitously captured in the recent structure of the malarial parasite *plasmodium falciparum* transporter *Pf*HT1^20^. Nonetheless, *Pf*HT1 is very distantly related to GLUT proteins (Fig. S1), and while GLUT proteins show strict sugar specificity, *Pf*HT1 transports a wide range of different sugars, making it unclear whether this occluded state would constitute a good representative of occluded state in GLUT proteins ^20,23^. Somewhat unexpectedly, the glucose coordinating residues in *Pf*HT1 were found to be almost identical to those in *human* GLUT3 ^20,24,25^. Based on the position of TM7b and biochemical analysis, it was concluded that the extracellular gating helix had evolved to transport many sugars, rather than the sugar-binding site itself ^20^. Simply put, it was proposed that *Pf*HT1 was less selective in what sugars it transports as its extracellular gate shuts more easily. Whilst the allosteric coupling between TM7b and the sugar-binding pocket might be more pronounced in *Pf*HT1, we hypothesized that the fundamental basis for sugar-coupling should be conserved in the GLUT proteins ^8,20^. Here, using enhanced sampling molecular dynamic simulations and GLUT5-proteoliposome transport assays, we have reconstructed the GLUT5 transport cycle, deciphering the molecular determinants for fructose binding and extracellular TM7b gating.

## Results and discussion

### Modeling rat GLUT5 in all conformational states

To piece together the “rocker-switch” alternating-access mechanism for the GLUT proteins ^26^, we must correctly assemble the relevant conformational states along the transport pathway. We thus selected to focus our efforts on the fructose transporter GLUT5 for two reasons. Firstly, GLUT5 is the only GLUT protein with structures determined in both outward-open and inward-open conformations, which principal component analysis of n = 17 sugar porter structures, confirmed represents the two end states ^20^. Secondly, how non-glucose sugars are recognized by GLUT proteins is unknown, and a computational framework for a fructose-specific transporter would help to understand substrate specificity more broadly.

Initially, to fill in the “missing” GLUT5 conformational states, homology models of *rat* GLUT5 were generated using relevant structures as templates: outward-occluded (*human* GLUT3), occluded (*Pf*HT1) and inward-occluded (*E. coli* XylE) and inward-open *bison* GLUT5 (Methods). To assess the stability of the generated homology models, hundreds of nanoseconds-long MD simulations on each of these models were performed in a model POPC membrane bilayer (Methods, Table 1). Each model was stable during the simulation, with a slightly higher RMSD in fully-occluded, inward-occluded, and inward-open models, likely reflective of their intrinsic dynamics in absence of substrate as well as ICH mobility when the N-and C-terminal bundles are no longer held together by salt bridges (Fig. S2). Overall, we concluded that the *rat* GLUT5 models had reached an acceptable dynamic equilibrium.

**Table 1.** GLUT5 model and simulation details.

| State modeled | Type of model | ICH5 resolved? | Template structure (if applicable) | PDB code | Percentage identity to rat GLUT5 | Simulation time (ns) |
| --- | --- | --- | --- | --- | --- | --- |
| Outward open | Structure | Yes |  | 4ybq | 100% | 523 |
| Outward occluded | Model | Yes | Human GLUT3 | 4zw9 | 40.8% | 298 |
| Fully occluded | Model | No | *Plasmodium falciparum* hexose transporter (PfHT1) | 6rw3 | 26.0% | 381 |
| Inward occluded | Model | No | *E. Coli* xylose transporter (XylE) | 4ja3 | 23.3% | 158 |
| Inward open | Model | No | Bovine GLUT5 | 4yb9 | 76.7% | 550 |

During substrate translocation by MFS transporters, cavity-closing contacts are predominantly formed between TM1 and TM7 on the outside (extracellular gate) and between TM4 and TM10 on the inside (intracellular gate) ^8^ (Fig. 1C). We can therefore monitor the distances between the centers of mass of the residues forming the extracellular gate (EG) and intracellular gate (IG) as a proxy for the conformational states sampled during simulations (Fig. 2A). As seen Fig. 2B, although the most populated gating distances deviate from the starting GLUT5 models (shown as filled-circles), all states equilibrated with mostly overlapping distributions. Notably, the largest deviation from the starting template is for the GLUT5 outward-occluded state modelled from *human* GLUT3, which we attribute to the fact the extracellular gate TM7b was stabilized in the crystal structure by the crystallization lipid monoolein ^15^. The non-filled gap between the occluded and inward-occluded state distributions corresponds to the larger global “rocker-switch” rearrangements (Fig. 2A), which are inaccessible over these short hundreds of ns-long time scales.

**Fig 2.**
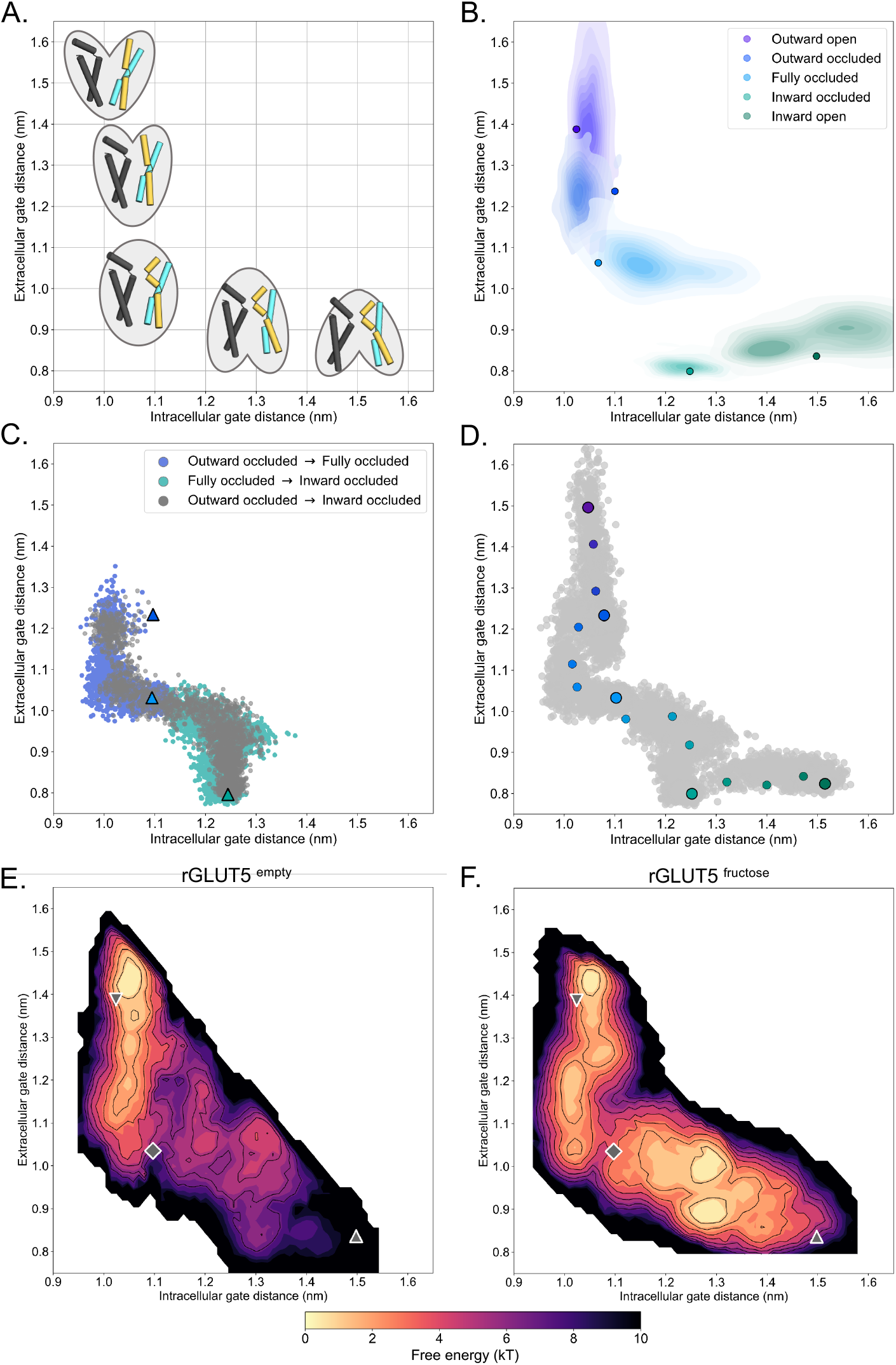
A free-energy landscape for D-fructose influx by GLUT5. **A**. A graphical illustration of the five major states of sugar porters, with the intracellular (IC) and extracellular (EC) gate distances on the x-axis and y-axis, respectively. Only TM1, TM4, TM7, and TM10 are drawn here, with the intent to highlight only major elements of the rocker switch conformational change. **B**. IC and EC gate population densities of atomistic simulations of each rGLUT5 homology model. Filled circles represent the starting configurations from each rGLUT5 homology model. **C**. Targeted MD (TMD) with bound fructose. Individual states are shown in triangles, with the following color schemes: outward occluded: deep blue, occluded: light blue, inward occluded: green. Grey dots represent all frames corresponding to TMD of outward occluded to inward occluded, skipping the occluded state. This follows a pathway similar to sequential TMD from outward occluded to fully occluded (blue circles), and fully occluded to inward occluded (teal circles). rGLUT5^empty^ TMD results can be found in Figure S3A. **D**. Beads chosen for the string simulations from TMD projected onto the space defined by the IC and EC gate distances for rGLUT5^fructose^. The cloud of grey dots represent all gate distance configurations through the TMD simulations, larger colored dots represent each initial homology model, and the smaller colored dots represent the beads between each of these models, which were chosen for the first iteration of the string-of-swarms method. Beads for rGLUT5^empty^ found in Fig. S3B. **E**. Free energy surface for rGLUT5^empty^. The triangles and diamond illustrate the gate measurements for the following homology models: outward open (top left triangle), fully occluded (diamond), and inward open (bottom right). **F**. Free energy surface for rGLUT5^fructose^, with the same homology model projection as in E.

### Interpolating between models of states using targeted MD simulations

To fully sample the conformational space between the occluded and inward-occluded GLUT5 states, enhanced sampling simulations are necessary. As a first step, we chose here targeted MD (TMD) simulations to interpolate between all five major states ^27^, applying a moving harmonic potential restraint to all heavy atoms in GLUT5. Given the uncertainty of the *Pf*HT1 occluded state as a suitable model in GLUT proteins, we performed TMD from the outward-occluded to the inward-occluded state either via the occluded state model or directly between these two states, both with and without fructose present. In both these targeted MD simulation protocols, we find that GLUT5 passes through a conformation in which the positioning of the gates closely matched the ones in the occluded model based on *Pf*HT1 (Fig. 2C, Fig. S3A).

Having confirmed that the *Pf*HT1 structure is a reasonable approximation for the occluded state in GLUT5, we aimed to characterize the most probable transition pathway linking the outward-open and inward-open states, and calculate the free energy surface lining this pathway. To this end, we used the string-of-swarms method for GLUT5 both in its apo (rGLUT5^empty^) and fructose-bound (rGLUT5^fructose^) conditions ^28^. In brief, each of the five different structural models were represented as beads along a string, with a further eleven beads added from configurations extracted from the TMD, yielding in total 16 beads spanning a tentative initial pathway defined in terms of their intracellular and extracellular gate distance (Fig. 2D, Fig. S3B). From each of these beads, many short trajectories were launched (swarms) to iteratively seek an energy minimum along the string path (see Methods). In this approach, the string simulations converge when the string diffuses around an equilibrium position. This protocol has proven effective to sufficiently explore computational space for complex conformational changes ^28^. After ∼100 iterations, the strings had converged (Fig. S3C), indicating that an equilibrium position was found. Nevertheless, we continued to run another ∼450 to 650 iterations to ensure exhaustive sampling of the entire transition pathway, enabling an appropriate estimation of the free energy along the converged path.

### The free-energy landscape of GLUT5 with and without fructose bound

Once the strings had converged and equilibrium was reached, we calculated free energy surfaces (FES) based on the transitions of all equilibrium swarm simulations (see Methods). Comparing the free energy surfaces between these two conditions reveals obvious differences between rGLUT5^empty^ and rGLUT5^fructose^ simulations (Fig. 2E, F). In the absence of fructose, the outward-open state is the most energetically favorable, with higher energy barriers to either occluded or inward-facing states (Fig. 2E). This calculation is consistent with experimental observations for the related XylE that show that the outward-facing state is the most populated in the absence of sugar ^29^. The free energy surface of GLUT5^empty^ is also consistent with structures that have shown that the strictly-conserved salt-bridge network is only present on the cytoplasmic inside ^8^, stabilizing the outward-facing state. Single point mutations to the salt-bridge network residues have indeed been shown to arrest GLUT transporters in the inward-facing conformation ^16,30^. In the presence of sugar, the inward-facing states become accessible and are of similar energetic stability to the outward-facing states (Fig. 2F). The heights of the free energy barriers between outward and inward-facing states in presence and absence of substrate, respectively, are consistent with measurements GLUT kinetics, as rates have been shown to be 100-fold faster for substrate-bound than for empty-occluded transitions ^2,22^. In other words, we can directly see the effect of substrate binding on sugar-induced conformational changes. Based on these overall differences matching experimental observations, we conclude the free energy landscape represents the physiologically-relevant GLUT5 conformational cycle.

In the presence of fructose, the occluded state model of GLUT5 is exactly positioned between the two energetically-favorable outward- and inward-facing states (Fig. 2F). Consistent with a transition state, the occluded state model is located on the highest energetic barrier along the lowest energy pathway between the two opposite-facing conformations. Moreover, transition into the occluded state from the outward states is energetically unfavorable for GLUT5 without sugar, but the presence of fructose clearly lowers the activation barrier (Fig 2E, F). These calculations are also consistent with the fact that GLUT transporters are required to spontaneously reset to the opposite-facing conformation through an empty occluded transition ^8^, i.e., local gates rearrangements controlling intermediate states must be able to spontaneously close in the absence of sugar.

### Conformational stabilization of D-fructose coordination in the occluded state

Since the presence of D-fructose lowers the energetic barriers between outward- and inward-facing states (Fig. 2F), we reasoned that we should be able to extract the molecular determinants for D-fructose coordination in the occluded state from these simulations. Based on extensive biochemical data and the glucose-bound *human* GLUT3 structure, we know that D-glucose is transported with the C1-OH group facing the bottom of the cavity (endofacial) and the C6-OH group facing the top (exofacial) ^8,31-33^. It is expected that D-fructose will be likewise transported by GLUT5 with the C1-OH group facing the endofacial direction, since substituents to fructose were better tolerated when added to the C6-OH position ^34,35^. D-fructose was unconstrained during TMD and string simulations. To evaluate the conformational heterogeneity of D-fructose, we binned the energy landscape, extracted configurations corresponding to each bin, and then clustered the fructose poses for these ensembles of configurations (see Methods). As seen in Fig. 3A, in the outward-open and outward-occluded conformations, D-fructose does not display any preferential binding mode, and the C1-OH group has no preferential orientation (brown sphere). In contrast, in the occluded state, the sugar becomes highly coordinated, adopting a single well-defined binding pose in approximately 65% of conformations that is 9-fold more populated than the next most abundant pose (Fig. 3A, B, Methods Table 3). Remarkably, the two most preferred poses are very similar to the orientation that both D-glucose and D-xylose have adopted in previously determined crystal structures (Fig. 3C)^8,15,17^.

**Fig 3.**
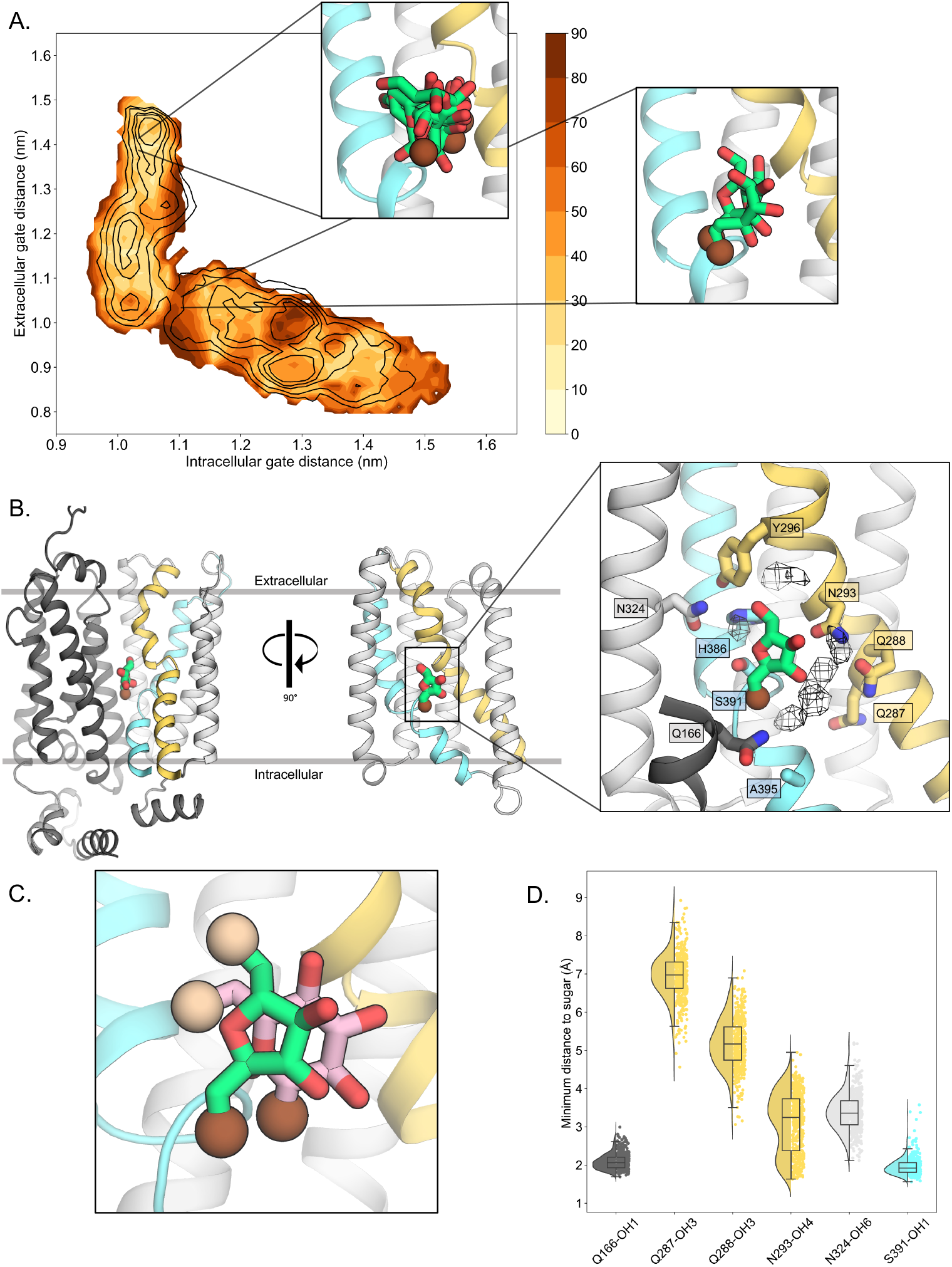
D-fructose becomes highly-coordinated in the occluded conformation. **A**. Sugar coordination of the fructose-bound simulations superimposed onto the free energy landscape for rGLUT5^fructose^, colored according to the frequency of the most populated cluster in each bin, therefore darker colors indicate a more consistent pose (see Methods for bin description). Snapshots extracted from two bins, corresponding to the outward open or occluded states respectively, depicting ∼70% of total pose variability (see Table 3 in Methods) are shown as inserts. GLUT5 helices, D-fructose, and C1 hydroxyl group shown as in panel B. **B**. An overview of a representative binding pose of fructose in occluded GLUT5, colored as in Figure 1. The D-fructose C1 hydroxyl group is colored as a brown sphere for orientation purposes. Selected interacting residues shown as sticks, and the density of waters near the fructose in the occluded state is shown as a black mesh (see Methods). **C**. The coordination of fructose is oriented the same as previously determined glucose positions, such as in *Pf*HT1 (PDB:6rw3, pink). The C6 hydroxyl group is shown as an enlarged sphere colored tan, and the C1 hydroxyl group in brown as in panel B. The fructose pose chosen here is the most populated cluster, as seen in Methods, Table 3. **D**. The distribution of distances of indicated fructose hydroxyl groups to certain side chains. Distances shown are calculated from the most populated cluster in the occluded state bin (Table 3, Methods).

Upon closer inspection of the fructose-bound state, we see that the serine residue S391, located between the TM10a-b breakpoint, coordinates with the C1-OH group of fructose (Fig. 3B, D), which is a glycine residue in the glucose-specific GLUT members (Fig. S1). The mutation of S391 to alanine in rat GLUT5 has been shown to weaken D-fructose binding ^14^. Consistent with D-glucose bound sugar porter structures^15,17,20,24,36^, the strictly-conserved TM5 glutamine Q166 is also forming tight hydrogen-bond interactions to the C1-OH group (Fig. 3B, D), which is a critical interaction required for coupling the N- and C-terminal bundles *i*.*e*., Q166 is the only residue on the N-terminal bundle that has been found to coordinate the substrate sugar in D-glucose bound sugar porter structures^8,15^. Unexpectedly, the TM7a glutamine residues Q287 and Q288 coordinating the C2- and C3-OH groups of D-glucose in GLUT3 and other members (Fig. S1)^8^, are not forming direct hydrogen bonds with D-fructose in the GLUT5 simulations (Fig. 3B, D). We find, however, that the most highly coordinated waters in the MD simulations are located between Q287 and Q288 and D-fructose, which are within hydrogen-bond distance to the C2-OH and C3-OH groups (Fig. 3B). In GLUT1, GLUT3, GLUT4 structures the bulky TM10 tryptophan residue directly coordinates the C1-OH group of D-glucose^15,16,37^. GLUT5, however, lacks the TM10 tryptophan, and this residue is replaced by Ala395 (Fig. 3B). It appears that Ser391 in TM10 is instead forming the C1-OH hydrogen-bond interaction, which was unexpected. The interaction between the TM10 serine 391 and C1-OH means that Q287 side-chain would be too far away to directly coordinate the C2-OH group. Although this would mean the C2-OH is poorly coordinated, this reasoning is consistent with biochemical analysis. Specifically, GLUT5 can transport 2,5-dihydromannitol with similar kinetics to D-fructose, which is a sugar that is identical to the furanose form of D-fructose. but lacks the C2-OH group^35^. In contrast, D-fructose epimers that differ by the orientation in any of the other -OH groups are unable to bind, since they do not show any cold competition for ^14^C-D-fructose uptake by *human* GLUT5 ^35^.

Upon superimposition of all the major conformations along the GLUT transport cycle, the TM7b asparagine was shown to be the only sugar-coordinating residue significantly changing its position during the transport cycle^20^. Because the TM7b asparagine residue is strictly conserved in all GLUT transporters and related sugar porters, it was proposed that the recruitment of the TM7b asparagine is a key and generic interaction required for coupling sugar binding with extracellular TM7b gating^8^. Consistently, in the simulations, the TM7b asparagine (N293) is also well positioned to coordinate the C4-OH group of D-fructose in the fully occluded state, and generally maintains hydrogen-bond distance (Fig. 3B, D). In addition to the TM7b asparagine (N293), a TM7b gating tyrosine (Y296) also forms an interaction to a histidine residue (H386) in TM10a (Fig. 4A). Both the TM10a histidine and the TM7b tyrosine are unique to GLUT5 (Fig. S1) ^14^ and GLUT5 variants H386F, H386A, and Y296A have been shown to severely diminish D-fructose binding ^14^. The TM7b tyrosine appears to also interact with an asparagine residue (N324) (Fig. 4A), which is also generally within hydrogen bond distance to the C6-OH group of D-fructose (Fig. 3D). As such, the TM7b gate seems to be connected both indirectly and directly to the sugar-binding site.

**Fig 4.**
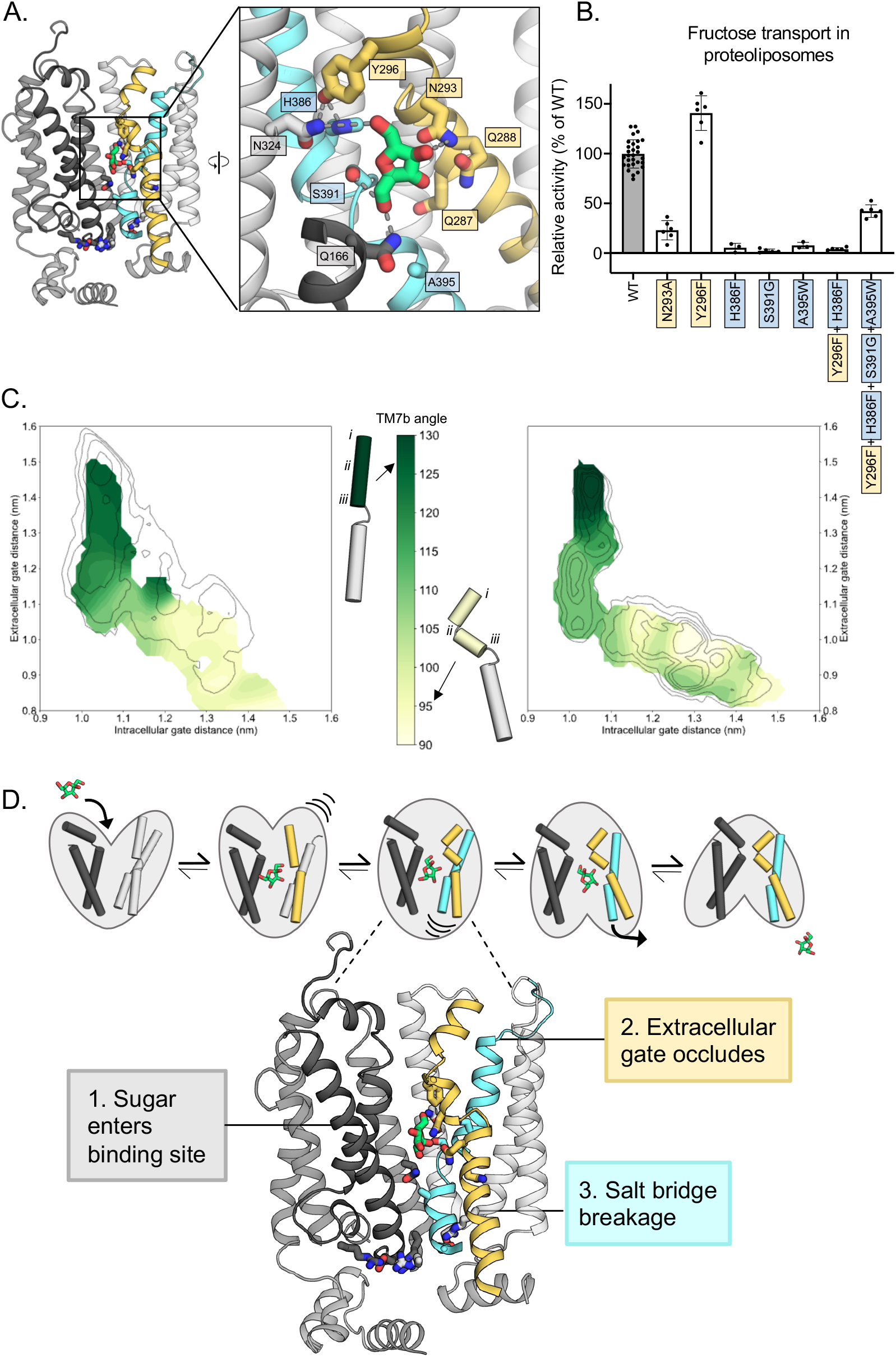
Confirming the residues required for coupling sugar binding to the extracellular gate TM7b and the intracellular gate TM10b. **A**. An overview of the residues connecting the coupling of TM7b breakage during sugar binding to the movement of TM10b. The occluded state is represented here. When N293 is pulled towards the fructose hydroxyl group on C4, TM7b is kinked, and Y296 is rotated towards TM10. Y296 is then able to interact with H386 on TM10a. Colouring and objects shown as described in Figure 1. TM2, TM11, and ICH4 are omitted for clarity, though the salt bridge residue R407 on TM11 remains. The inlay highlights key residue interactions with the sugar, the Y296-H386 interactions, and the Y296-N324 interactions. Dashes represent possible hydrogen bond interactions (as shown in Fig. 3B and 3D, not including Y296-H386 or Y296-N324). **B**. ^14^C D-fructose transport activity of WT rGLUT5 and mutations related to the coupling of fructose binding with TM7 breakage and TM10 movement, residue names are coloured as in panel A. Uptake of ^14^C-D-fructose (black bars) in proteoliposomes of rat GLUT5 variants relative to WT. Errors bars represent s.e.m. of at least 6 independent measurements taken from two to three independent liposome reconstitutions. **C**. TM7b angle for rGLUT5^empty^ (left) and rGLUT5^fructose^(right) superimposed onto the respective free energy landscapes. This angle is calculated by measuring the vector formed between residue groups *i,ii,iii* as shown on the protein cartoons (see Methods). The angle in the outward open state is approximately 130 degrees, and will bend to nearly a 90 degree angle towards the inward-facing states. **D**. Our updated view on the molecular determinates of sugar-induced conformational change in GLUT5. Residues moving to interact with the sugar in the binding site induce a conformational change in TM7b (yellow), occluding the extracellular gate. This breakage of TM7b in a fully occluded state is communicated to an inward opening via TM10b (cyan). Interactions between TM7b (Y296) to TM10a (H386) and TM8(N324), as well as between the fructose and S391, stabilize TM10a. The TM10b, which is separated from TM10a by the GPXPXP motif, is uncoupled from TM10a, and will move away from TM4b. This movement will then induce breakage of the salt bridge residues to an inward open state. Coloring as in Figure 1.

To strengthen the proposed MD model for D-fructose coordination, we evaluated ^14^C-D-fructose transport of purified GLUT5 variants reconstituted into liposomes using our recently developed transport assay^38^ (see Methods for details). We focused the mutagenesis on GLUT residues that are different in GLUT5 from GLUT transporters incapable of D-fructose transport^39^. Consistently, the H386F and S391G variants that were constructed to match the most common residue in the other GLUT transporters ^15^, abolished D-fructose transport (Fig. 4B). As shown recently^38^, the A395W variant is also incapable of D-fructose transport (Fig. 4B). All GLUT5 variants were unable to transport D-glucose (Fig. S4C). The H386F and S391G mutations are particularly informative, as these residues do not coordinate D-glucose in the glucose-bound GLUT structures^15,16,37^. Confirming the importance of the strictly-conserved TM7b asparagine N293, its substitution to alanine also severely reduced transport (Fig.4B).

### Coupling between fructose binding and TM7b gating rearrangements

In the *Pf*HT1 occluded structure, TM7b was found to have broken into an elbow-shaped helix, with close contacts to TM1 at the break-point. In all inward-facing states, the gating helix TM7b remains at a sharp angle ^8^. Structural information suggests that the stabilization of TM7b *via* the asparagine coordination to a substrate sugar would induce TM7b to transition from a bent to a broken-helix conformation ^8,20^. To assess this structural transition, we compared the angle formed by TM7b throughout both rGLUT5^empty^ and rGLUT5^fructose^ simulations (Fig. 4C). Consistently, we observe that TM7b comparatively forms a sharper angle earlier in the transport cycle when sugar is present, indicating that indeed the conformational state of TM7b is connected with sugar recognition, and suggesting a mechanism whereby D-fructose binding induces transition into the occluded states. The angle of TM7b further decreases upon transition into the occluded state to fully shut the outside gate.

GLUT transporters harbor two strictly-conserved tyrosine residues in TM7b^14^, which form the substrate-occlusion in the outward-occluded conformation^8^. In *Pf*HT1, TM7b residues are considerably more polar and the TM7b tyrosine residues have been replaced by serine and asparagine^20^. *Pf*HT1 mutations of TM7b gating residues were found to be just as critical to transport as residues coordinating D-glucose^20^. Furthermore, TM7b variants retaining transport competence shifted the substrate preference of *Pf*HT1 from D-glucose towards D-fructose^8,20^. It was, thus, proposed that *Pf*HT1 had not evolved the sugar binding site to be able to transport many sugars, but its extracellular gate. In MD simulations of GLUT5, the TM7b tyrosine residue Y296 that formed an interaction with H386, is located next to these occlusion-forming Y297 and Y298 residues. In most other GLUT isoforms, Y296 is a phenylalanine, and so to assess this interaction further we measured transport of an Y296F variant. Somewhat surprisingly, the Y296F variant retained robust ^14^C-D-fructose transport with an ∼40% improved performance (k_cat_/*K*_M_) (Fig. 4B, S4A). Upon closer inspection, we find that the TM7b tyrosine is a phenylalanine residue in around 30% of more distantly-related GLUT5 homologues (Fig. S5). Indeed, independent co-evolution analysis corroborates that Y298 and H383 residues are forming an interaction in the occluded state of GLUT5^40^. Moreover, in yeast-based forward-evolution screens of hexose transporters, the equivalent residue to Y298 in TM7b was the only singe-residue-variant uncovered with a shifted sugar preference from D-glucose towards D-xylose^41,42^. It is possible that the divergence of the TM7b residue from phenylalanine to tyrosine has evolved so that GLUT5 can transport an as yet, unidentified sugar in some tissues, which has been speculated previously, since GLUT5 is expressed in tissues with very low circulating levels of D-fructose^5^.

To further demonstrate the importance of coupling between TM7b and the sugar binding site, we combined single point mutations that individually abolished D-fructose transport (H285F, S391G, A395W) with the TM7b variant (Y296F). Interestingly, the combined variant is now re-able to transport D-fructose transport with ∼40% wildtype activity (Fig. 4B, S4B), although it is still unable to transport D-glucose (Fig. S4C). Thus, while the GLUT5 binding site has evolved to specifically coordinate D-fructose, there is plasticity in the sugar binding pocket that can allow for alternative interactions when combined with differences to extracellular TM7b gating residues *i*.*e*., as was recently demonstrated in the malarial parasite transporter *Pf*HT1^20^.

Extracellular TM7b gate closure in the occluded conformation must somehow trigger the breakage of the intracellular salt-bridge network on the inside in order for the two bundles to come apart. The mutational analysis of sugar binding residues supports the modelled D-fructose, which implies that that the substrate-sugar stabilizes both the closure of TM7b, but and also an interaction with TM10a. MD simulations shows that when TM10a becomes locked in place by interaction with the substrate sugar, TM10b is able to move more independently (Fig. S6A,B), which is facilitated by a very mobile GPXPXP helix-break motif (Fig. S1, S5). In the simulations, we see that the TM7b angle decreases from about 140 degrees to about 115 degrees without any noticeable change in TM10b (Fig. S6A). However, as the TM7b angle reaches 115 degrees in the occluded state, TM10b undergoes a large shift in position. Interestingly, the salt-bridge residues, particularly those located between TM4 and TM11, do not fully break apart until TM10b has finished rearranging (Fig. S6B, S6C). This would indeed be consistent with the coordinated coupling between the inward movement of TM7b triggering the outward movement of TM10b to break the inter-bundle salt-bridge network.

## Discussion

GLUT transporters are often presented as text-book examples of how small molecule transporters are functional equivalents of soluble enzymes. Yet, despite extensive kinetic, biochemical and physiological analysis, we have a poor understanding of how GLUT structures fit into such a molecular description. Here, for the first time, we can confirm that the occluded state structure of *Pf*HT1 ^20^ provides a suitable template for modelling the transition state in a GLUT transporter. The classical description of enzyme catalysis is that there is relatively weak binding of the substrate to the enzyme in the initial state, but a tight binding in the transition state ^43,44^. This conceptual framework implies that in GLUT proteins the sugar would bind more tightly to the transition state, which would be consistent the Induced Transition Fit of transport catalysis proposed by Klingenberg ref. ^44^. More specifically, in the occluded state, we find that TM7b is broken over the sugar-binding site to better coordinate D-fructose. The fundamental difference between enzymes and transporters is that the structure of the transition state determines the activation barrier for global conformational changes in transporters, whereas in enzymes, the barrier is imposed by substrate remodeling in the transition state ^44^. Here, we indeed observe that the energy barrier for conversion between states is clearly lowered by the coordination of D-fructose. It is worth mentioning, that we have focused on the role of the sugar coupling in influx rather than efflux, because the affinities for D-glucose are reported to be 10-fold lower on the inside in other GLUT transporters and homologues ^8,45^ and salt-bridge formation between the two bundles was more difficult to model (see Methods, Fig. S6D).

By measuring GLUT1 kinetics at different temperatures, an activation barrier (Ea) of around 10 kcal/mol has been reported ^22^. This relatively low activation barrier roughly corresponds to the breakage of a few salt-bridges, which matches the expectation for the intracellular salt-bridge-rich GLUTs. The D-glucose binding energies has been estimated to be around 9 kcal/mol for GLUT3 ^46^, which is consistent with sugar binding required to generate the global transitions by inducing formation of the occluded state. Although the transition state represents the highest energetic barriers between opposite-facing conformations in MD simulation of GLUT5, the height of the activation barrier cannot be reliably calculated from our simulations for several different reasons. Firstly, the energy barriers are estimated along a path that describes structural transitions in the extracellular and intracellular gates, rather than all conformational changes across the entire protein. Moreover, our models consider a membrane bilayer made from POPC lipids, whereas it is well established that transport by GLUT proteins requires the presence of anionic lipids. The fact that the activation barrier for GLUT1 has been shown to increase from 10 to 16 kcal/mol in liposomes made from lipids with longer fatty acids highlights just how sensitive GLUT proteins are to the lipid composition ^47^. Here, we chose to use a neutral lipid composition to avoid complications related to the anticipated timescales needed to equilibrate a complex bilayer.

The coordination of D-fructose in the occluded state is consistent with previous biochemical analysis, the coordination of D-glucose, and the GLUT5 proteoliposome assays carried out here. The asparagine residue located in the beginning of TM7b is conserved across the entire sugar porter superfamily, and is the only sugar coordinating residue significantly changing its position during the transport cycle^8,20^. Our simulations show that N293 is recruited to coordinate D-fructose in a similar manner to D-glucose, and we propose that the TM7b asparagine is a generic interaction required for coupling the binding of any transported monosaccharide sugar with the extracellular TM7b gate.

Once TM7b has been stabilized by a substrate sugar *via* N293, the closure of TM7b is influenced by residues along the length of TM7b. In the promiscuous sugar transporter *Pf*HT1, the strictly-conserved TM7b gating tyrosine residues found in GLUT1-14, have been replaced by serine and asparagine (Figure S1) ^20^. These more polar residues enable closing of the outside gate more easily and play a role in catalyzing transport of different sugars. In contrast, GLUT5 is a highly-specific sugar transporter and has a finely-tuned extracellular TM7b gate. More specifically, upon TM7b gate closure, the tyrosine residue Y298 preceding the “YY” motif forms a unique pairing to a histidine residue peripheral to the sugar-binding site. The histidine can interact both with the TM7b tyrosine as well as hydrogen bond to an asparagine residue interacting with fructose. It is poignant that we observe a connection between the sugar-binding site and the TM7b tyrosine residue, which is located at the region wherein the TM7b helix transitions from a bent to a broken helix in the occluded state. The importance of TM7b was also observed in the *E. coli* xylose symporter XylE. Whilst XylE binds D-glucose in the same manner and with the same affinity as in *human* GLUT3, the XylE protein is incapable of transporting the sugar, i.e., D-glucose is a dead-end inhibitor ^48^. However, the mutation of the residue corresponding to the TM7b tyrosine in XylE (L297F) together with a sugar binding site mutant (Q175L), enables XylE to transport D-glucose while retaining 75% of wild-type D-xylose transport ^49^. Illustrating the importance of the TM7b gate, we have been able to reconstruct a GLUT5 sugar binding site with individually “dead” mutants that can recover D-fructose transport when combined with the mutagenesis of an TM7b gating residue. Taken together, our work strengthens the proposal that TM7b should be considered as an extension of the sugar-binding site ^20^.

## Conclusions

We conclude the molecular determinants for sugar transport are an intricate coupling between an extracellular gate, a sugar-binding site, and an intracellular salt-bridge network (Fig. 4D). Weakly binding sugars are able to induce large conformational changes in GLUT proteins by conformational stabilization of a transition state that can already be spontaneously populated. Rather than a substrate binding site that achieves strict specificity by having a high-affinity for the substrate, GLUT proteins have allosterically coupled sugar binding with an extracellular gate that forms the high-affinity transition-state instead. Presumably, this substrate-coupling pathway ensures that sugar-binding does not become rate-limiting, and so enables GLUT proteins to catalyse fast sugar flux at physiological relevant blood-glucose concentrations in the mM range. The recent type 2 diabetes drug empagliflozin in complex with the sodium-coupled glucose transporter SGLT2, demonstrates how selective inhibition was achieved by the aglycone of the glucoside inhibitor interacting with the mobile TM1a-b and TM6a-b half-helices ^50^. In many aspects, while GLUT proteins are referred to as rocker-switch proteins, their asymmetric binding mode gives rise to gating elements closely resembling the structural transitions seen in rocking-bundle proteins, like SGLT2 ^8^. Such an intricate coupling indicates that pharmacological control of GLUT proteins might best be accomplished by small molecules targeting gating regions extending from the sugar binding site.

## Materials and Methods

### Molecular dynamics simulations

#### Protein modeling and atomistic simulations

Residue numbing for rGLUT5 is based on the UNIPROT entry of rGLUT5: P43427, all generated models begin at residue E7 and end at residue V480. The starting models for rGLUT5 in each state were generated using homology modeling with MODELLER version 10.1 ^51^. A summary of the details of these models and subsequent simulations are found in Table 1. The unresolved TM1-TM2 loop from chain A of the outward-open structure of rGLUT5, PDB:4ybq, was modeled using MODELLER. Human GLUT3, 4zw9 ^25^, served as template for the rGLUT5 outward-occluded model. *Pf*HT1 PDB:6rw3 ^20^ chain C served as template for the rGLUT5 fully-occluded model. XylE PDB:4ja3 ^18^, served as template for the rGLUT5 inward-occluded model. Bovine GLUT5, PDB:4yb9 ^14^, served as template for the rGLUT5 inward-open model. Intracellular helix 5 (ICH5) is not present or incomplete in several structures (see Table 1) and therefore the rGLUT5 ICH5 (residues M457-V480) was added to the sequence alignment for homology modeling as a template.

Each rGLUT5 protein model was placed into a POPC bilayer with ∼122 lipids on the top leaflet, and ∼124 on the bottom, and solvated in a water box with 150mM NaCl using CHARMM-GUI ^52^. The total box size before equilibration was 10×10×11nm. All parameters of the system were described using CHARMM36m.

Each system underwent energy minimization using steepest descent, followed by system equilibration for a total of 187.5ps where positional restraints on the protein and POPC lipids were gradually released. Production MD was then run using 2fs timesteps in GROMACS version 2019.1 ^53^. Temperature was maintained at 303.15K using Nose-Hoover temperature coupling, using three separate groups for protein, lipid bilayer, and the solvent. Pressure was maintained at 1bar using the Parrinello-Rahman barostat with semiisotropic coupling, using a time constant of 5ps and a compressibility of 4.5 × 10^−5^ bar^−1^. Hydrogen bonds were constrained using LINCS ^54^, electrostatic interactions modelled with a 1.2nm cutoff, while long-range electrostatics were calculated with particle mesh Ewald (PME). Simulation length can be found in Table 1.

#### Targeted molecular dynamics

Targeted MD (TMD) was performed in a stepwise fashion between states to ensure that the initial string would cover the determined sugar porter conformational space observed thus far. Four main TMD protocols were used: rGLUT5^empty^ Outward open - Inward open, rGLUT5^empty^ Inward open - Outward open, rGLUT5^fructose^ Outward open - Inward open, and rGLUT5^fructose^ Inward open - Outward open.

TMD was performed using GROMACS version 2019.5 patched with PLUMED version 2.5.5 ^55^. Each TMD condition was performed in a stepwise, iterative fashion. The first TMD run of each of the four conditions was performed biasing stepwise, either the outward open structure or inward open homology model towards the respective targets, the outward occluded or inward occluded models. All structures and models used, be it as a starting point or as a target for the TMD, are from unequilibrated (not from the aforementioned MD) structures/models, to ensure that the TMD was not generated from a local minima distant from a desired state. The positions of all heavy atoms were biased in a geometric space with incrementally increasing harmonic restraints, initially starting at 0 kJ/mol/nm and increasing to 2500 kJ/mol/nm at step 5000. After 5000 steps, the force applied was squared every 150000 steps until the heavy atom RMSD of the system was within about 0.05nm of their target conformation. After this was achieved, the final frame of the TMD run was used to generate the next TMD run’s input model for each condition. Table 2 details each TMD run length, starting and ending conformations for each condition, and the RMSD of the final TMD timepoint.

**Table 2.** Summary of targeted MD simulations.

| rGLUT5<sup>empty</sup> Outward open - Inward open (condition 1) |  |  |  |  |
| --- | --- | --- | --- | --- |
| TMD number | Starting configuration | Target state | Time (ps) | Final heavy atom RMSD (nm) |
| 1.1 | Out open structure | Outward occluded | 11,220 | 0.048415 |
| 1.2 | TMD 1.1 final frame | Fully occluded | 10,890 | 0.050022 |
| 1.3 | TMD 1.2 final frame | Inward occluded | 8,120 | 0.062960 |
| 1.4 | TMD 1.3 final frame | Inward open | 10,780 | 0.052656 |
| 1.5 (skipping occluded state validation) | TMD 1.1 final frame | Inward occluded | 10,440 | 0.051464 |
| rGLUT5<sup>empty</sup> Inward open - Outward open (condition 2) |  |  |  |  |
| TMD number | Starting configuration | Target state | Time (ps) | Final heavy atom RMSD (nm) |
| 2.1 | In open homology model | Inward occluded | 9,620 | 0.066614 |
| 2.2 | TMD 2.1 final frame | Fully occluded | 8,620 | 0.053146 |
| 2.3 | TMD 2.2 final frame | Outward occluded | 7,760 | 0.058147 |
| 2.4 | TMD 2.3 final frame | Outward open | 9,120 | 0.054561 |
| 2.5 (skipping occluded state validation) | TMD 2.1 final frame | Outward occluded | 11,840 | 0.048901 |
| rGLUT5<sup>fructose</sup> Outward open - Inward open (condition 3) |  |  |  |  |
| TMD number | Starting configuration | Target state | Time (ps) | Final heavy atom RMSD (nm) |
| 3.1 | Out open structure with fructose bound | Outward occluded | 8,500 | 0.065881 |
| 3.2 | TMD 3.1 final frame | Fully occluded | 10,780 | 0.052316 |
| 3.3 | TMD 3.2 final frame | Inward occluded | 12,630 | 0.059851 |
| 3.4 | TMD 3.3 final frame | Inward open | 10,260 | 0.045643 |
| 3.5 (skipping occluded state validation) | TMD 3.1 final frame | Inward occluded | 12,480 | 0.055612 |
| rGLUT5<sup>fructose</sup> Inward open - Outward open (condition 4) |  |  |  |  |
| TMD number | Starting configuration | Target state | Time (ps) | Final heavy atom RMSD (nm) |
| 4.1 | In open homology model with fructose bound | Inward occluded | 10,000 | 0.075750 |
| 4.2 | TMD 4.1 final frame | Fully occluded | 12,980 | 0.048203 |
| 4.3 | TMD 4.2 final frame | Outward occluded | 11,780 | 0.049837 |
| 4.4 | TMD 4.3 final frame | Outward open | 12,420 | 0.049001 |
| 4.5 (skipping occluded state validation) | TMD 4.1 final frame | Outward occluded | 12,480 | 0.049699 |

**Table 3.** Analysis of sugar binding pose in outward open and occluded states.

| Bin closest to: **Outward open state**<br>Total frames in bin: **1000**<br>Total possible clusters for sugar pose: **217** |  |  | Bin closest to: **Occluded state**<br>Total frames in bin: **690**<br>Total possible clusters for sugar pose: **116** |  |  |
| --- | --- | --- | --- | --- | --- |
| Cluster number | Number of frames in cluster | Percentage of total frames in bin | Cluster number | Number of frames in cluster | Percentage of total frames in bin |
| 1 | 234 | 23.40% | 1 | 447 | 64.78% |
| 2 | 123 | 12.30% | 2 | 52 | 7.54% |
| 3 | 106 | 10.60% | 3 | 24 | 3.48% |
| 4 | 86 | 8.60% | 4 | 23 | 3.33% |
| 5 | 59 | 5.90% | 5,6,7 | 3 | 0.43% |
| 6 | 38 | 3.80% | 8-12 | 2 | 0.29% |
| 7 | 36 | 3.60% | 13-116 | 1 | 0.14% |
| 8-12 | 32-10 | 3.20% - 1.00% |  |  |  |
| 13-16 | 5-2 | 0.50% - 0.20% |  |  |  |
| 17-217 | 1 | 0.10% |  |  |  |

For rGLUT5 ^fructose^ TMD runs, beta-D-fructofuranose was placed in the outward open structure or inward open model based on the positioning of glucose in hGLUT1 (4pyp)^15^ after structural alignment with the models in PyMol version 2.5.0. The fructose-bound outward open structure or inward open model were then briefly energy minimized to ensure no sugar and water atoms clashing during simulation. During TMD runs, fructose coordinates were left unbiased.

#### Limitations of inward open to outward open simulations in regards to salt bridge distances

Initially, as described above, TMD was also performed with both rGLUT5^empty^ and rGLUT5^fructose^, from inward open to outward open. However, string simulations performed of these conditions did not converge. Upon examination of features of these simulations, we could see the state-dependent salt-bridge residues losing contact in states where they should not (Fig. S6D), and thus we elected to focus further simulations on GLUT5 influx, as discussed in the main text.

#### Collective variables selection

Collective variables (CVs) were chosen based on features that were transferable to other sugar porters, and that would separate different functional states. Two CVs were used for this state differentiation, measuring opening of the extracellular and the intracellular gates, respectively (Fig. 1C, 2A, S1). The distance between the centers of mass of the extracellular gating parts of the transmembrane helices TM1 (residues 36 to 43) and TM7 (residues 295 to 301) were used to measure opening of the extracellular gate (referred to as extracellular gate distance). The distance between the centers of mass of the intracellular gating parts of the transmembrane helices TM4 (residues 142 to 151) and TM10 (residues 392 to 400) were used to measure opening of the intracellular gate (referred to as intracellular gate distance).

#### String preparation

For each of the conditions, snapshots corresponding to points lining the string were extracted from the TMD runs (referred to as beads hereafter). 16 beads were chosen in total, five of which correspond to the outward open, outward occluded, fully occluded, inward occluded, and inward open models, and based on the first or final frames of the TMD runs. The other 11 beads were chosen to cover uniformly the CV space between states (Fig. 2D for rGLUT5^fructose^, Fig. S3B for rGLUT5^empry^).

#### String method with swarms of trajectories

The string simulations with swarms of trajectories were performed as described in ref ^28^, with a brief summary as follows. With the exception of bead 0 and bead 15 (outward open and inward open models), which were held fixed and therefore not simulated in each run, each bead along the string underwent several simulation steps in every iteration of the string simulations. <u>Step 1: short string reparametrization and CV equilibration</u>. The CV values were extracted for each bead, and the relevant system was equilibrated with a 10,000 kJ nm ^−2^ harmonic force potential acting on each CV for 30ps. <u>Step 2: swarms of trajectories</u>. From each bead, 32 swarms were launched in parallel and run for 10ps each. <u>Step 3: calculate CV drift for next iteration</u>. The drift per bead was calculated by measuring the average of the CV distance over the simulation swarm. <u>Step 4</u>: Using the updated CV coordinates, the string was reparametrized so that the beads were equidistantly placed along the string, therefore stopping each bead from falling into nearby energy minima. Details of this reparameterization can be found in (29). Then, the iteration was complete and the next iteration could begin, with the initial simulation restraining the system in the reparametrized CV space. The rGLUT5 ^fructose^ simulations were run for a total of 552 iterations, and the rGLUT5 ^empty^ simulations were run for a total of 745 iterations.

The code for running the string simulations with the conditions above, as well as a tutorial and simple system setup and analysis code can be found on GitHub at https://github.com/delemottelab/string-method-swarms-trajectories. All simulation parameters of the string simulations were the same as mentioned above, with the exception of GROMACS version (2020.5 instead of 2019.1), and the use of a V-rescale thermostat instead of Nose-Hoover.

#### Free energy landscape calculation

The free energy landscapes as depicted in Fig. 2E and 2F were calculated from the transition matrix of the swarm simulations in the CV space once they were determined to be in equilibrium (Fig. S3C), after about 100 iterations. Therefore, 452 iterations of data were used to calculate rGLUT5 ^fructose^ free energy surfaces, and 645 iterations of data for rGLUT5 ^empty^. First, a time-lagged independent component analysis (TICA) of the CVs was performed over every iteration, for each bead, using each start and end position in the swarm, to efficiently separate the data for state discretization. Next, these TICA projections were clustered using k-means clustering. Finally, a Markov state model (MSM) was constructed with n=100 clusters and a kernel density estimation (KDE) of the resulting MSM was projected into the 2D collective variable space, using a bandwidth of 0.05, on a 55×55 grid.

#### Analysis of simulations

The resulting free energy landscape is defined on a 55×55 grid. For the analysis of protein features (such as sugar coordination, TM7b angle, and salt bridge distances), each of the bins in the grid was analyzed independently. For this, structural snapshots from the endpoints of swarm simulations corresponding to the CV values of each bin were extracted, with a maximum of 1000 frames per bin (see Fig. S7A).

In the sugar coordination analysis, the snapshots extracted for each bin were aligned on the entire protein position in cartesian space. Then, clustering was performed on each of the sugar coordinates using gmx cluster with the Jarvis-Patrick algorithm and a cutoff of 0.08nm for each cluster center. The percentage of total frames occupied by the most populated cluster is presented in Fig. 3A. Table 3 summarizes the clusters for two representative bins, the outward open and occluded state. The area shaded grey in the table indicate clusters used in the Fig. 3A inserts, highlighting the most populated clusters summing to ∼70% of the total possible sugar poses in a bin. The most populated cluster for the occluded state, highlighted with a red border, is shown in Fig. 3C. Distances between certain sugar hydroxyl groups and residue side chains, as shown in Fig. 3D, are calculated from this cluster as well. These measurements are the minimum closest distance between any atom of a given hydroxyl group, to any atom of a residue side chain for each frame in this bin (n=447).

In other feature analysis such as TM7b angle or salt bridge distances, the snapshots extracted for each bin were analyzed in Python version 3.8.5 using MD Analysis version 2.0.0^12^ and plotted in matplotlib version 3.3.4.

Images overlaying simulation features with an energy surface (such as Fig. 3A and Fig. 4C) use an abstraction of the free energy surfaces as depicted in Fig 2E and 2F. A depiction of this abstraction can be found in Fig. S7B.

To estimate average properties for each grid point (*X*_*i*_), weighted averages (*W*) are reported, using the weights for each snapshot estimated from the MSM (*w*_*i*_).

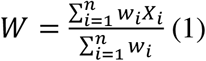

The TM7b angle *θ* was calculated as the angle between two vectors defined by two groups of residues center of mass (COM):

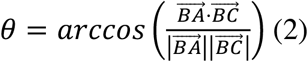

where A represents the vector of positions of residues 289-291 COM, B the vector of positions of residues 296-298 COM and C the vector of positions of residues 304-306 COM.

The TM10b RMSD was calculated as the weighted average of each grid’s RMSD of the backbone of residues 391-401 after superposition of all backbone atoms onto the outward open structure 4yb9.

The state-dependent salt bridge residue distance is calculated as the minimum distance averaged between residue pairs E151-R407, and E400-R158.

The water density as seen in Fig. 3B was generated from a trajectory of snapshots from the most occupied sugar pose cluster of the occluded state (red outline, Table 3). The density was calculated in VMD using the VolMap tool, measuring average water density in all frames, of waters within 5Å of the fructose molecule. The resulting density is visualized in PyMol, with an isomesh density cutoff of 0.7.

The aforementioned Python codes written for all analysis and free energy landscape calculations can be found at: https://github.com/semccomas/GLUT5_string

#### rGLUT5 homolog detection and sequence logo generation

The rGLUT5 sequence was searched using NCBI pBLAST against a non-redundant protein database, with 1000 maximum target sequences and E-value threshold of 0.05. The sequences were then aligned with ClustalOmega (https://www.ebi.ac.uk/Tools/msa/clustalo/). From this alignment, a sequence logo was generated with Weblogo3 (https://weblogo.threeplusone.com) locally.

### Functional activity of rGLUT5 mutants

#### Construct design and cloning

The full-length sequence of rGLUT5 (Uniprot: P43427) was used for functional assays, with 2 alterations to the sequence. The deglycosylation mutation N50Y is present, and several C-terminal residues are retained after TEV cleavage. Both are underlined in the following sequence. rGLUT5 wild-type (WT): AMEKEDQEKTGKLTLVLALATFLAAFGSSFQYGYNVAAVNSPSEFMQQFY<u>Y</u>D TYYDRNKENIESFTLTLLWSLTVSMFPFGGFIGSLMVGFLVNNLGRKGALLFNNIFSILPAILMGCSKIAKSFEIIIASRLLVGICAGISSNVVPMYLGELAPKNLRGALGVVPQLFITVGILVAQLFGLRSVLASEEGWPILLGLTGVPAGLQLLLLPFFPESPRYLLIQKKNESAAEKALQTLRGWKDVDMEMEEIRKEDEAEKAAGFISVWKLFRMQSLRWQLISTIVLMAGQQLSGVNAIYYYADQIYLSAGVKSNDVQYVTA GTGAVNVFMTMVTVFVVELWGRRNLLLIGFSTCLTACIVLTVALALQNTISWMPYVSIVCVIVYVIGHAVGPSPIPALFITEIFLQSSRPSAYMIGGSVHWLSNFIVGLIFPFIQVGLGPYSFIIFAIICLLTTIYIFMVVPETKGRTFVEINQIFAKKNKVSDVYP EKEEKELNDLPPATREQ<u>ENLYFQ.</u>

#### GLUT5 *overexpression and membrane isolation*

The sequence (WT or with desired mutations) was cloned into the GAL-inducible vector pDDGFP2, containing a His_8_ sequence and TEV cleavage site^56,57^. Mutants were generated by overlap PCR. The vector was cloned into the *Saccharomyces cerevisiae* strain FGY217. rGLUT5 was overexpressed and subsequently purified as previously described^56,58^. Briefly, 12L of *S. cerevisiae* cells were grown in -URA medium containing 0.1% (w/v) D-glucose. Cells were incubated at 30°C, inducing with 2% (w/v, final) D-galactose once OD_600_ reached 0.6. 22h after induction, cells were harvested, resuspended in buffer containing 50mM Tris-HCl (pH 7.6), 1mM EDTA, and 600mM sorbitol, and lysed mechanically. Unlysed cells and debris was removed by centrifugation at 4°C and 10,000g for 10m, then membranes were then isolated by ultracentrifugation at 4°C and 195,000g for 2h. Isolated membranes were then homogenized in 20mM Tris-HCl (pH 7.6), 300mM sucrose, and 0.1mM CaCl_2_.

#### Rat GLUT5 purification

Membranes were solubilized using 1% *n*-dodecyl-β-D-maltopyranoside (DDM, Glycon) in equilibration buffer containing 1x PBS, 150mM NaCl, and 10% (w/v) glycerol for 1h. Unsolubilized membranes were removed by ultracentrifugation at 4°C and 195,000g for 45m. The remaining supernatant was incubated with 20mM imidazole and 15ml Ni-NTA resin (Qiagen) for 3h with mild agitation. This solution was then transferred to a 30ml gravity-flow column (Bio-Rad). Immobilized protein was then washed twice with a solution containing equilibration buffer, 0.1% DDM, and increasing concentrations of imidazole: 20mM then 30mM. 30ml of protein was eluted with equilibration buffer containing 250mM imidazole. The eluted sample was dialyzed overnight in a buffer of 20mM Tris-HCL (pH 7.6), 150mM NaCl, and 0.03% DDM, adding equimolar ratio of TEV to the dialysis bag. Dialyzed sample was passed through 5ml HisTrap columns (GE Healthcare) to isolate cleaved rGLUT5 protein, which was then concentrated (50KDa MWCO, Amicon) and injected to a Superose 6 column using a flow rate of 0.4 ml/min in a buffer of 20mM Tris-HCL (pH 7.6), 150mM NaCl, 0.03% DDM. Fractions corresponding to the monomer peak of GLUT5 were pooled (Fig. S4D) and concentrated to 2mg ml^−1^ before flash freezing aliquots of protein into liquid nitrogen.

#### GLUT5 Proteoliposome transport assay

rGLUT5 protein was incorporated into proteoliposomes and measured using a radiolabeled sugar as previously described for related sugar porters^20^. Briefly, liposomes were prepared in 500µl batches by mixing total bovine brain lipid extract (Sigma Aldrich, final 30mg ml^−1^) and cholesteryl-hemisuccinate powder (Sigma Aldrich, final 6mg ml^−1^) were mixed in a buffer of 10mM Tris-HCl (pH 7.6) and 2mM MgSO_4_. The lipid mix was then subjected to several rounds of flash-freeze and thawing cycles, and then sonication. Large lipid particles were removed by centrifugation at room temperature and 16,000g for 15 minutes. 20µg of purified rGLUT5 protein was added to 500µl of liposomes, and was flash-frozen, then thawed, and extruded through a 400nm filter (LiposoFast, Avestin).

For steady-state and time course measurement 20µl of prepared proteoliposomes were added to 2µl of [^14^C]D-fructose (6µM, American Radiolabelled Chemicals) and incubated for 2 minutes (time course measurement in 30 second intervals) at room temperature before stopping the reaction with 1ml of Tris-MgSO_4_ buffer, and filtering through a 0.22µm filter (Millipore), washing with 6ml Tris-MgSO_4_ buffer. Filters were transferred to scintillation vials, applying 5ml of scintillation liquid (Ultima Gold, Perkin Elmer) before measuring by scintillation counting. All reported measurements are repeated in triplicates, using empty liposomes as baseline noise, and WT activity at 100% activity.

For measuring glucose transport activity of rGLUT5 mutants, the same procedure as above is performed, using instead 2µl of [^14^C]D-glucose (6µM, American Radiolabelled Chemicals). As above, empty liposomes are used as baseline noise in the experiments, and WT activity for D-fructose transport is used 100% activity.

For the kinetic analysis of Y296F, 20µl of prepared proteoliposomes were diluted in 20µl of increasing concentrations of D-fructose (1mM – 30 mM, in Tris-MgSO_4_ buffer) with constant stochiometric amounts of [^14^C]D-fructose. The *K*_*M*_ and *V*_*max*_ values for D-fructose transport could be determined by measuring initial transport velocities at 60 s. The recorded decay counts of the transported D-fructose were baseline corrected with counts from empty liposomes and fitted to Michaelis–Menten kinetics using nonlinear regression by GraphPad Prism 9.0.

## Acknowledgements

We thank Sergio Perez Conesa for support using the string of swarms method and related discussions. We thank Albert Suades for providing purified protein of rGLUT5 mutant A395W. This work was funded by grants from Novo Nordisk foundation (to D.D.) and The Knut and Alice Wallenberg Foundation (to D.D.) and Göran Gustafsson foundation (to D.D and L.D). LD acknowledges SciLifeLab and the Swedish Research Council (VR 2019-02433) for funding. The MD simulations were performed on resources provided by the Swedish National Infrastructure for Computing (SNIC) on Beskow at the PDC Center for High Performance Computing (PDC-HPC).

## Author contribution

D.D. and L.D designed research; S.M performed molecular dynamics simulations; S.M, D.D and L.D analyzed data; S.M and T.R performed biochemical experiments; S.M, D.M, T.R, C.A, M.B, D.D and L.D wrote the paper. All authors approved the final manuscript.

## References

1 Holman, G. D. Structure, function and regulation of mammalian glucose transporters of the SLC2 family. Pflugers Arch 472, 1155–1175 (2020). https://doi.org:10.1007/s00424-020-02411-3

2 Mueckler, M. & Thorens, B. The SLC2 (GLUT) family of membrane transporters. Mol Aspects Med 34, 121–138 (2013). https://doi.org:10.1016/j.mam.2012.07.001

3 Thorens, B. GLUT2, glucose sensing and glucose homeostasis. Diabetologia 58, 221–232 (2015). https://doi.org:10.1007/s00125-014-3451-1

4 Huang, S. & Czech, M. P. The GLUT4 glucose transporter. Cell Metab 5, 237–252 (2007). https://doi.org:10.1016/j.cmet.2007.03.006

5 Douard, V. & Ferraris, R. P. Regulation of the fructose transporter GLUT5 in health and disease. Am J Physiol Endocrinol Metab 295, E227–237 (2008). https://doi.org:10.1152/ajpendo.90245.2008

6 Kayano, T. et al. Human facilitative glucose transporters. Isolation, functional characterization, and gene localization of cDNAs encoding an isoform (GLUT5) expressed in small intestine, kidney, muscle, and adipose tissue and an unusual glucose transporter pseudogene-like sequence (GLUT6). J Biol Chem 265, 13276–13282 (1990).

7 Koepsell, H. Glucose transporters in brain in health and disease. Pflugers Arch 472, 1299–1343 (2020). https://doi.org:10.1007/s00424-020-02441-x

8 Drew, D., North, R. A., Nagarathinam, K. & Tanabe, M. Structures and General Transport Mechanisms by the Major Facilitator Superfamily (MFS). Chem Rev (2021). https://doi.org:10.1021/acs.chemrev.0c00983

9 Zhang, Y., Zhang, Y., Sun, K., Meng, Z. & Chen, L. The SLC transporter in nutrient and metabolic sensing, regulation, and drug development. J Mol Cell Biol 11, 1–13 (2019). https://doi.org:10.1093/jmcb/mjy052

10 Ancey, P. B., Contat, C. & Meylan, E. Glucose transporters in cancer - from tumor cells to the tumor microenvironment. FEBS J 285, 2926–2943 (2018). https://doi.org:10.1111/febs.14577

11 Drew, D., North, R. A., Nagarathinam, K. & Tanabe, M. Structures and General Transport Mechanisms by the Major Facilitator Superfamily (MFS). Chem Rev 121, 5289–5335 (2021). https://doi.org:10.1021/acs.chemrev.0c00983

12 Pao, S. S., Paulsen, I. T. & Saier, M. H., Jr. Major facilitator superfamily. Microbiol Mol Biol Rev 62, 1–34 (1998).

13 Maiden, M. C., Davis, E. O., Baldwin, S. A., Moore, D. C. & Henderson, P. J. Mammalian and bacterial sugar transport proteins are homologous. Nature 325, 641–643 (1987). https://doi.org:10.1038/325641a0

14 Nomura, N. et al. Structure and mechanism of the mammalian fructose transporter GLUT5. Nature 526, 397–401 (2015). https://doi.org:10.1038/nature14909

15 Deng, D. et al. Molecular basis of ligand recognition and transport by glucose transporters. Nature 526, 391–396 (2015). https://doi.org:10.1038/nature14655

16 Deng, D. et al. Crystal structure of the human glucose transporter GLUT1. Nature 510, 121–125 (2014). https://doi.org:10.1038/nature13306

17 Sun, L. et al. Crystal structure of a bacterial homologue of glucose transporters GLUT1-4. Nature 490, 361–366 (2012). https://doi.org:10.1038/nature11524

18 Quistgaard, E. M., Low, C., Moberg, P., Tresaugues, L. & Nordlund, P. Structural basis for substrate transport in the GLUT-homology family of monosaccharide transporters. Nat Struct Mol Biol 20, 766–768 (2013). https://doi.org:10.1038/nsmb.2569

19 Wisedchaisri, G., Park, M. S., Iadanza, M. G., Zheng, H. & Gonen, T. Proton-coupled sugar transport in the prototypical major facilitator superfamily protein XylE. Nat Commun 5, 4521 (2014). https://doi.org:10.1038/ncomms5521

20 Qureshi, A. A. et al. The molecular basis for sugar import in malaria parasites. Nature 578, 321–325 (2020). https://doi.org:10.1038/s41586-020-1963-z

21 Paulsen, P. A., Custodio, T. F. & Pedersen, B. P. Crystal structure of the plant symporter STP10 illuminates sugar uptake mechanism in monosaccharide transporter superfamily. Nat Commun 10, 407 (2019). https://doi.org:10.1038/s41467-018-08176-9

22 Lowe, A. G. & Walmsley, A. R. The kinetics of glucose transport in human red blood cells. Biochim Biophys Acta 857, 146–154 (1986). https://doi.org:10.1016/0005-2736(86)90342-1

23 Woodrow, C. J., Penny, J. I. & Krishna, S. Intraerythrocytic Plasmodium falciparum expresses a high affinity facilitative hexose transporter. J Biol Chem 274, 7272–7277 (1999).

24 Jiang, X. et al. Structural Basis for Blocking Sugar Uptake into the Malaria Parasite Plasmodium falciparum. Cell 183, 258–268 e212 (2020). https://doi.org:10.1016/j.cell.2020.08.015

25 Deng, D. et al. Molecular basis of ligand recognition and transport by glucose transporters. Nature (2015). https://doi.org:10.1038/nature14655

26 Drew, D. & Boudker, O. Shared Molecular Mechanisms of Membrane Transporters. Annu Rev Biochem 85, 543–572 (2016). https://doi.org:10.1146/annurev-biochem-060815-014520

27 Schlitter, J., Engels, M. & Kruger, P. Targeted molecular dynamics: a new approach for searching pathways of conformational transitions. J Mol Graph 12, 84–89 (1994). https://doi.org:10.1016/0263-7855(94)80072-3

28 Fleetwood, O., Matricon, P., Carlsson, J. & Delemotte, L. Energy Landscapes Reveal Agonist Control of G Protein-Coupled Receptor Activation via Microswitches. Biochemistry 59, 880–891 (2020). https://doi.org:10.1021/acs.biochem.9b00842

29 Jia, R. et al. Hydrogen-deuterium exchange mass spectrometry captures distinct dynamics upon substrate and inhibitor binding to a transporter. Nat Commun 11, 6162 (2020). https://doi.org:10.1038/s41467-020-20032-3

30 Schurmann, A. et al. Role of conserved arginine and glutamate residues on the cytosolic surface of glucose transporters for transporter function. Biochemistry 36, 12897–12902 (1997). https://doi.org:10.1021/bi971173c

31 Holman, G. D. Chemical biology probes of mammalian GLUT structure and function. Biochem J 475, 3511–3534 (2018). https://doi.org:10.1042/BCJ20170677

32 Barnett, J. E., Holman, G. D. & Munday, K. A. Structural requirements for binding to the sugar-transport system of the human erythrocyte. Biochem J 131, 211–221 (1973). https://doi.org:10.1042/bj1310211

33 Barnett, J. E., Holman, G. D., Chalkley, R. A. & Munday, K. A. Evidence for two asymmetric conformational states in the human erythrocyte sugar-transport system. Biochem J 145, 417–429 (1975). https://doi.org:10.1042/bj1450417a

34 Tatibouet, A., Yang, J., Morin, C. & Holman, G. D. Synthesis and evaluation of fructose analogues as inhibitors of the D-fructose transporter GLUT5. Bioorg Med Chem 8, 1825–1833 (2000). https://doi.org:10.1016/s0968-0896(00)00108-5

35 Yang, J., Dowden, J., Tatibouet, A., Hatanaka, Y. & Holman, G. D. Development of high-affinity ligands and photoaffinity labels for the D-fructose transporter GLUT5. Biochem J 367, 533–539 (2002). https://doi.org:10.1042/BJ20020843

36 Bavnhoj, L., Paulsen, P. A., Flores-Canales, J. C., Schiott, B. & Pedersen, B. P. Molecular mechanism of sugar transport in plants unveiled by structures of glucose/H(+) symporter STP10. Nat Plants 7, 1409–1419 (2021). https://doi.org:10.1038/s41477-021-00992-0

37 Yuan, Y. et al. Cryo-EM structure of human glucose transporter GLUT4. Nat Commun 13, 2671 (2022). https://doi.org:10.1038/s41467-022-30235-5

38 Suades A, Qureshi A, McComas S.E, Coinçon M, Rudling A, Chatzikyriakidou Y, Landreh M, Carlsson J, Drew D. Establishing mammalian GLUT kinetics and lipid preferences in a reconstituted-liposome system. (2022). Research Square https://doi.org/10.21203/rs.3.rs-2176110/v1

39 Li, Q. et al. Cloning and functional characterization of the human GLUT7 isoform SLC2A7 from the small intestine. Am J Physiol Gastrointest Liver Physiol 287, G236–242 (2004). https://doi.org:10.1152/ajpgi.00396.2003

40 Mitrovic D M. S. E., Alleva C, Bonaccorsi M, Drew D, Delemotte L. Reconstructing the transport cycle in the sugar porter superfamily using coevolution-powered machine learning. bioRxiv (2022).: https://doi.org/10.1101/2022.09.24.509294

41 Wang, M., Yu, C. & Zhao, H. Identification of an important motif that controls the activity and specificity of sugar transporters. Biotechnol Bioeng 113, 1460–1467 (2016). https://doi.org:10.1002/bit.25926

42 Wang, M., Li, S. & Zhao, H. Design and engineering of intracellular-metabolite-sensing/regulation gene circuits in Saccharomyces cerevisiae. Biotechnol Bioeng 113, 206–215 (2016). https://doi.org:10.1002/bit.25676

43 Henzler-Wildman, K. & Kern, D. Dynamic personalities of proteins. Nature 450, 964–972 (2007). https://doi.org:10.1038/nature06522

44 Klingenberg, M. Ligand-protein interaction in biomembrane carriers. The induced transition fit of transport catalysis. Biochemistry 44, 8563–8570 (2005). https://doi.org:10.1021/bi050543r

45 Cloherty, E. K., Heard, K. S. & Carruthers, A. Human erythrocyte sugar transport is incompatible with available carrier models. Biochemistry 35, 10411–10421 (1996). https://doi.org:10.1021/bi953077m

46 Liang, H., Bourdon, A. K., Chen, L. Y., Phelix, C. F. & Perry, G. Gibbs Free-Energy Gradient along the Path of Glucose Transport through Human Glucose Transporter 3. ACS Chem Neurosci 9, 2815–2823 (2018). https://doi.org:10.1021/acschemneuro.8b00223

47 Carruthers, A. & Melchior, D. L. Human erythrocyte hexose transporter activity is governed by bilayer lipid composition in reconstituted vesicles. Biochemistry 23, 6901–6911 (1984). https://doi.org:10.1021/bi00321a096

48 Farwick, A., Bruder, S., Schadeweg, V., Oreb, M. & Boles, E. Engineering of yeast hexose transporters to transport D-xylose without inhibition by D-glucose. Proc Natl Acad Sci U S A 111, 5159–5164 (2014). https://doi.org:10.1073/pnas.1323464111

49 Madej, M. G., Sun, L., Yan, N. & Kaback, H. R. Functional architecture of MFS D-glucose transporters. Proc Natl Acad Sci U S A 111, E719–727 (2014). https://doi.org:10.1073/pnas.1400336111

50 Niu, Y. et al. Structural basis of inhibition of the human SGLT2-MAP17 glucose transporter. Nature 601, 280–284 (2022). https://doi.org:10.1038/s41586-021-04212-9

51 Sali, A. & Blundell, T. L. Comparative protein modelling by satisfaction of spatial restraints. J Mol Biol 234, 779–815 (1993). https://doi.org:10.1006/jmbi.1993.1626

52 Jo, S., Kim, T., Iyer, V. G. & Im, W. CHARMM-GUI: a web-based graphical user interface for CHARMM. J Comput Chem 29, 1859–1865 (2008). https://doi.org:10.1002/jcc.20945

53 Abraham, M. M. T; Roland, S; Páll, S; Smith, J; Hess, B; Lindahl, E. GROMACS: High performance molecular simulations through multi-level parallelism from laptops to supercomputers. SoftwareX 1, 19–25 (2015).

54 Hess, B. P-LINCS: A parallel linear constraint solver for molecular simulation. Journal of Chemical Theory and Computation 4, 116–122 (2008).

55 Tribello, G. A., Bonomi, M., Branduardi, D., Camilloni, C. & Bussi, G. PLUMED 2: New feathers for an old bird. Comput. Phys. Commun. 185, 604–613 (2014).

56 Drew, D. et al. GFP-based optimization scheme for the overexpression and purification of eukaryotic membrane proteins in Saccharomyces cerevisiae. Nat Protoc 3, 784–798 (2008). https://doi.org:10.1038/nprot.2008.44

57 Newstead, S., Kim, H., von Heijne, G., Iwata, S. & Drew, D. High-throughput fluorescent-based optimization of eukaryotic membrane protein overexpression and purification in Saccharomyces cerevisiae. Proc Natl Acad Sci U S A 104, 13936–13941 (2007).

