## Supplementary Material for "Determinants of sugar-induced influx in the mammalian fructose transporter GLUT5"

### Figure S1

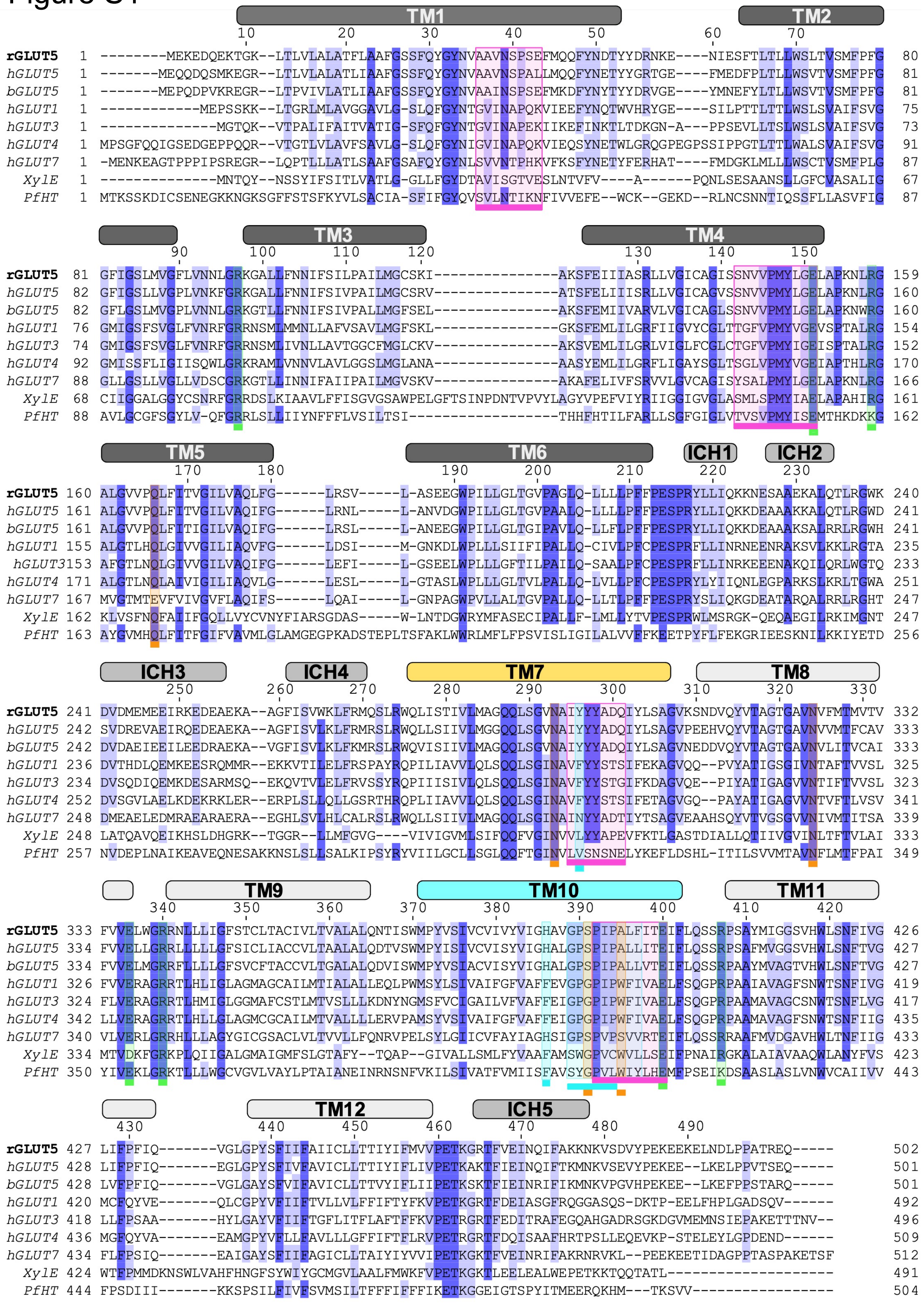
TM1
TM2
10 20 30 40 50 60 70
rGLUT5 1 --------MEKEDQEKTGK--LTLVLALATFLAAFGSSFQYGYNVAAVNSPSEFMQQFYNDTYYDRNKE----NIESFTLTLLWSLTVSMFPFG 80
hGLUT5 1 -------MEQQDQSMKEGR--LTLVLALATLIAAFGSSFQYGYNVAAVNSPALLMQQFYNETYYGRTGE----FMEDFPLTLLWSVTVSMFPFG 81
bGLUT5 1 -------MEPQDPVKREGR--LTPVIVLATLIAAFGSSFQYGYNVAAINSPSEFMKDFYNYTYYDRVGE----YMNEFYLTLLWSVTVSMFPFG 81
hGLUT1 1 ------------MEPSSKK--LTGRLMLAVGGAVLG-SLQFGYNTGVINAPQKVIEEFYNQTWVHRYGE----SILPTTLTTLWSLSVAIFSVG 75
hGLUT3 1 --------------MGTQK--VTPALIFAITVATIG-SFQFGYNTGVINAPEKIIKEFINKTLTDKGN-A---PPSEVLLTSLWSLSVAIFSVG 73
hGLUT4 1 MPSGFQQIGSEDGEPPQQR--VTGTLVLAVFSAVLG-SLQFGYNIGVINAPQKVIEQSYNETWLGRQGPEGPSSIPPGTLTTLWALSVAIFSVG 91
hGLUT7 1 -MENKEAGTPPPIPSREGR--LQPTLLLATLSAAFGSAFQYGYNLSVVNTPHKVFKSFYNETYFERHAT----FMDGKLMLLLWSCTVSMFPLG 87
XylE 1 --------------MNTQY--NSSYIFSITLVATLG-GLLFGYDTAVISGTVESLNTVFV----A------PQNLSESAANSLLGFCVASALIG 67
PfHT 1 MTKSSKDICSENEGKKNGKSGFFSTSFKYVLSACIA-SFIFGYQVSVLNTIKNFIVVEFE--WCK--GEKD--RLNCSNNTIQSSFLLASVFIG 87
TM3
TM4
90 100 110 120 130 140 150
rGLUT5 81 GFIGSLMVGFLVNNLGRKGALLFNNIFSILPAILMGCSKI---------------AKSFEIIIASRLLVGICAGISSNVVPMYLGELAPKNLRG 159
hGLUT5 82 GFIGSLLVGPLVNKFGRKGALLFNNIFSIVPAILMGCSRV---------------ATSFELIIISRLLVGICAGVSSNVVPMYLGELAPKNLRG 160
bGLUT5 82 GFLGSLMVGPLVNNLGRKGTLLFNNIFSIVPALLMGFSEL---------------AKSFEMIIVARVLVGICAGLSSNVVPMYLGELAPKNWRG 160
hGLUT1 76 GMIGSFSVGLFVNRFGRRNSMLMMNLLAFVSAVLMGFSKL---------------GKSFEMLILGRFIIGVYCGLTTGFVPMYVGEVSPTALRG 154
hGLUT3 74 GMIGSFSVGLFVNRFGRRNSMLIVNLLAVTGGCFMGLCKV---------------AKSVEMLILGRLVIGLFCGLCTGFVPMYIGEISPTALRG 152
hGLUT4 92 GMISSFLIGIISQWLGRKRAMLVNNVLAVLGGSLMGLANA---------------AASYEMLILGRFLIGAYSGLTSGLVPMYVGEIAPTHLRG 170
hGLUT7 88 GLLGSLLVGLLVDSCGRKGTLLINNIFAIIPAILMGVSKV---------------AKAFELIVFSRVVLGVCAGISYSALPMYLGELAPKNLRG 166
XylE 68 CIIGGALGGYCSNRFGRRDSLKIAAVLFFISGVGSAWPELGFTSINPDNTVPVYLAGYVPEFVIYRIIGGIGVGLASMLSPMYIAELAPAHIRG 161
PfHT 88 AVLGCGFSGYLV-QFGRRLSLLIIYNFFFLVSILTSI-----------------THHFHTILFARLLSGFGIGLVTVSVPMYISEMTHKDKKG 162
TM5
TM6
ICH1
ICH2
170 180 190 200 210 220 230
rGLUT5 160 ALGVVPQLFITVGILVAQLFG------LRSV-----L-ASEEGWPILLGLTGVPAGLQ-LLLLPFFPESPRYLLIQKKNESAAEKALQTLRGWK 240
hGLUT5 161 ALGVVPQLFITVGILVAQIFG------LRNL-----L-ANVDGWPILLGLTGVPAALQ-LLLLPFFPESPRYLLIQKKDEAAAKKALQTLRGWD 241
bGLUT5 161 ALGVVPQLFITIGILVAQIFG------LRSL-----L-ANEEGWPILLGLTGIPAVLQ-LLFLPFFPESPRYLLIQKKDEAAAKSALRRLRGWH 241
hGLUT1 155 ALGTLHQLGIVVGILIAQVFG------LDSI-----M-GNKDLWPLLLSIIFIPALLQ-CIVLPFCPESPRFLLINRNEENRAKSVLKKLRGTA 235
hGLUT3 153 AFGTLNQLGIVVGILVAQIFG------LEFI-----L-GSEELWPLLLGFTILPAILQ-SAALPFCPESPRFLLINRKEEENAKQILQRLWGTQ 233
hGLUT4 171 ALGTLNQLAIVIGILIAQVLG------LESL-----L-GTASLWPLLLGLTVLPALLQ-LVLLPFCPESPRYLYIIQNLEGPARKSLKRLTGWA 251
hGLUT7 167 MVGTMTEVFVIVGVFLAQIFS------LQAI-----L-GNPAGWPVLLALTGVPALLQ-LLTLPFFPESPRYSLIQKGDEATARQALRRLRGHT 247
XylE 162 KLVSFNQFAIIFGQLLVYCVNYFIARSGDAS-----W-LNTDGWRYMFASECIPALLF-LMLLYTVPESPRWLMSRGK-QEQAEGILRKIMGNT 247
PfHT 163 AYGVMHQLFITFGIFVAVMLGLAMGEGPKADSTEPLTSFAKLWWRLMFLFPSVISLIGILALVVFFKEETPYFLFEKGRIEESKNILKKIYETD 256
ICH3
ICH4
TM7
TM8
250 260 270 280 290 300 310 320 330
rGLUT5 241 DVDMEMEEIRKEDEAEKA--AGFISVWKLFRMQSLRWQLISTIVLMAGQQLSGVNAIYYYADQIYLSAGVKSNDVQYVTAGTGAVNVFMTMVTV 332
hGLUT5 242 SVDREVAEIRQEDEAEKA--AGFISVLKLFRMRSLRWQLLSIIVLMGGQQLSGVNAIYYYADQIYLSAGVPEEHVQYVTAGTGAVNVVMTFCAV 333
bGLUT5 242 DVDAEIEEILEEDRAEKA--VGFISVLKLFKMRSLRWQVISIIVLMAGQQLSGVNAIYYYADQIYLSAGVNEDDVQYVTAGTGAVNVLITVCAI 333
hGLUT1 236 DVTHDLQEMKEESRQMMR--EKKVTILELFRSPAYRQPILIAVVLQLSQQLSGINAVFYYSTSIFEKAGVQQ--PVYATIGSGIVNTAFTVVSL 325
hGLUT3 234 DVSQDIQEMKDESARMSQ--EKQVTVLELFRVSSYRQPIIISIVLQLSQQLSGINAVFYYSTGIFKDAGVQE--PIYATIGAGVVNTIFTVVSL 323
hGLUT4 252 DVSGVLAELKDEKRKLER--ERPLSLLQLLGSRTHRQPLIIAVVLQLSQQLSGINAVFYYSTSIFETAGVGQ--PAYATIGAGVVNTVFTLVSV 341
hGLUT7 248 DMEAELEDMRAEARAERA--EGHLSVLHLCALRSLRWQLLSIIVLMAGQQLSGINAINYYADTIYTSAGVEAAHSQYVTVGSGVVNIVMTITSA 339
XylE 248 LATQAVQEIKHSLDHGRK--TGGR--LLMFGVG----VIVIGVMLSIFQQFVGINVVLYYAPEVFKTLGASTDIALLQTIIVGVINLTFTVLAI 333
PfHT 257 NVDEPLNAIKEAVEQNESAKKNSLSLLSALKIPSYRYVIILGCLLSGLQQFTGINVLVSNSNELYKEFLDSHL-ITILSVVMTAVNFLMTFPAI 349
TM9
TM10
TM11
340 350 360 370 380 390 400 410 420
rGLUT5 333 FVVELWGRRNLLLIGFSTCLTACIVLTVALALQNTISWMPYVSIVCVIVYVIGHAVGPSPIPALFITEIFLQSSRPSAYMIGGSVHWLSNFIVG 426
hGLUT5 334 FVVELLGRRLLLLLGFSICLIACCVLTAALALQDTVSWMPYISIVCVISYVIGHALGPSPIPALLITEIFLQSSRPSAFMVGGSVHWLSNFTVG 427
bGLUT5 334 FVVELMGRRFLLLLGFSVCFTACCVLTGALALQDVISWMPYVSIACVISYVIGHALGPSPIPALLVTEIFLQSSRPAAYMVAGTVHWLSNFTVG 427
hGLUT1 326 FVVERAGRRTLHLIGLAGMAGCAILMTIALALLEQLPWMSYLSIVAIFGFVAFFEVGPGPIPWFIVAELFSQGPRPAAIAVAGFSNWTSNFIVG 419
hGLUT3 324 FLVERAGRRTLHMIGLGGMAFCSTLMTVSLLLKDNYNGMSFVCIGAILVFVAFFEIGPGPIPWFIVAELFSQGPRPAAMAVAGCSNWTSNFLVG 417
hGLUT4 342 LLVERAGRRTLHLLGLAGMCGCAILMTVALLLLERVPAMSYVSIVAIFGFVAFFEIGPGPIPWFIVAELFSQGPRPAAMAVAGFSNWTSNFIIG 435
hGLUT7 340 VLVERLGRRHLLLAGYGICGSACLVLTVVLLFQNRVPELSYLGIICVFAYIAGHSIGPSPVPSVVRTEIFLQSSRRAAFMVDGAVHWLTNFIIG 433
XylE 334 MTVDKFGRKPLQIIGALGMAIGMFSLGTAFY--TQAP--GIVALLSMLFYVAAFAMSWGPVCWVLLSEIFPNAIRGKALAIAVAAQWLANYFVS 423
PfHT 350 YIVEKLGRKTLLLWGCVGVLVAYLPTAIANEINRNSNFVKILSIVATFVMIISFAVSYGPVLWIYLHEMFPSEIKDSAASLASLVNWVCAIIVV 443
TM12
ICH5
430 440 450 460 470 480 490
rGLUT5 427 LIFPFIQ-------VGLGPYSFIIFAIICLLTTIYIFMVVPETKGRTFVEINQIFAKKNKVSDVYPEKEEKELNDLPPATREQ----- 502
hGLUT5 428 LIFPFIQ-------EGLGPYSFIVFAVICLLTTIYIFLIVPETKAKTFIEINQIFTKMNKVSEVYPEKEE--LKELPPVTSEQ----- 501
bGLUT5 428 LVFPFIQ-------VGLGAYSFVIFAVICLLTTVYIFLIIPETKSKTFIEINRIFIKMNKVPGVHPEKEE--LKEFPPSTARQ----- 501
hGLUT1 420 MCFQYVE-------QLCGPYVFIIFTVLLVLFFIFTYFKVPETKGRTFDEIASGFRQGGASQS-DKTP-EELFHPLGADSQV------ 492
hGLUT3 418 LLFPSAA-------HYLGAYVFIIFTGFLITFLAFTFFKVPETRGRTFEDITRAFEGQAHGADRSGKDGVMEMNSIEPAKETTTNV-- 496
hGLUT4 436 MGFQYVA-------EAMGPYVFLLFAVLLLGFFIFTFLRVPETRGRTFDQISAAFHRTPSLLEQEVKP-STELEYLGPDEND------ 509
hGLUT7 434 FLFPSIQ-------EAIGAYSFIIFAGICLLTAIYIYVVIPETKGKTFVEINRIFAKRNRVKL--PEEKEETIDAGPPTASPAKETSF 512
XylE 424 WTFPMMDKNSWLVAHFHNGFSYWIYGCMGVLAALFMWKFVPETKGKTLEELEALWEPETKKTQQTATL-------------------- 491
PfHT 444 FPSDIII-------KKSPSILFIVFSVMSILTFFFIFFFIKETKGGEIGTSPYITMEERQKHM---TKSVV---------------- 504

**Figure S1. Sequence alignment** of *rat* GLUT5 to *human* and *bovine* GLUT5, as well as other GLUTs of interest and sugar porters used in homology modeling (GLUT3, PfHT1, and XylE). A dark blue color indicates >88% (8/9 matching) sequence identity, and a lighter blue color indicates a >77% (7/9 matching) sequence identity. The transmembrane and ICH helices are indicated by labelled squares above the alignment, colored as in Figure 1. The intracellular (on TM4 and TM10) and extracellular gates (on TM1 and TM7) are colored in pink squares. The boxes below sugar binding residues are colored in orange, residues involved in coupling of binding to conformational change as well as the GPXPXP motif in turquoise, and salt bridge residues in green. Residues with multiple features (such as on TM10) have stacked boxes.

Figure S2

A.

B.

C.

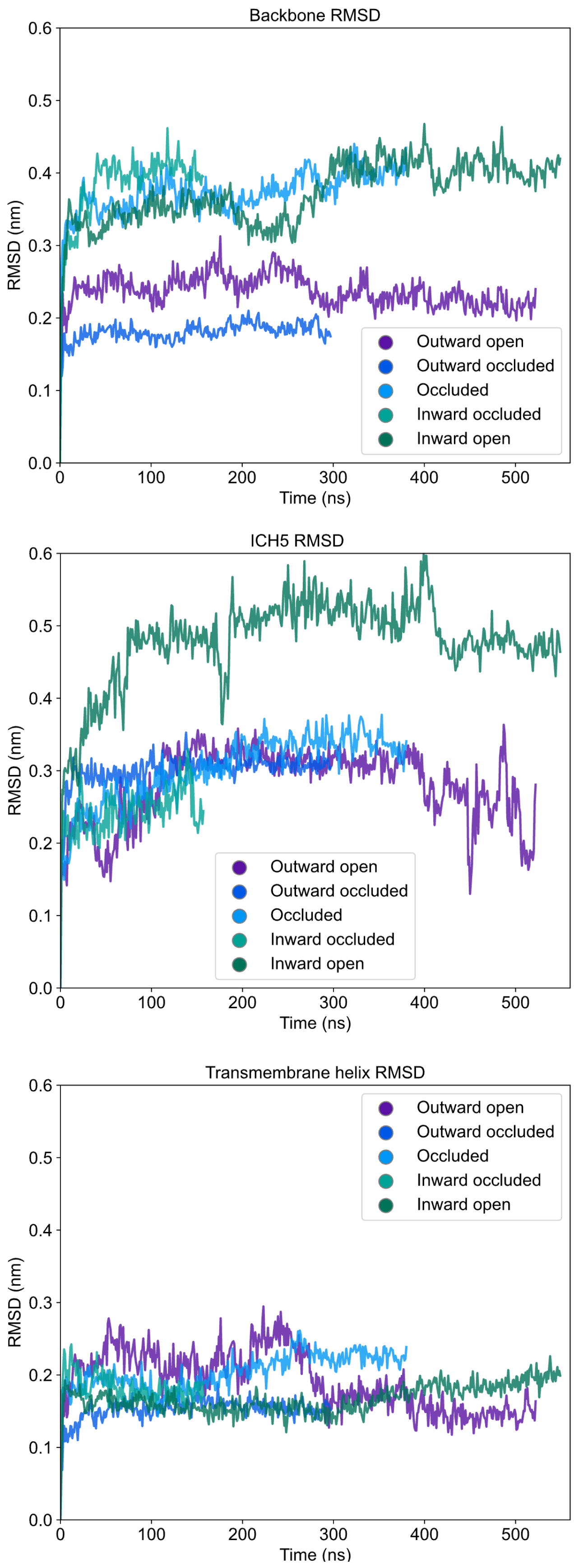

**Figure S2. Root mean squared deviation (RMSD) of atomistic simulations of rGLUT5 homology models. A.** Backbone RMSD calculated by aligning on all C-alpha atoms of the protein upon the original homology model, and reporting their RMSD. **B.** RMSD of ICH5 residues after self-aligning. **C.** RMSD of transmembrane helices 1-12 (as indicated in Fig. S1) after self-aligning.

### Figure S3

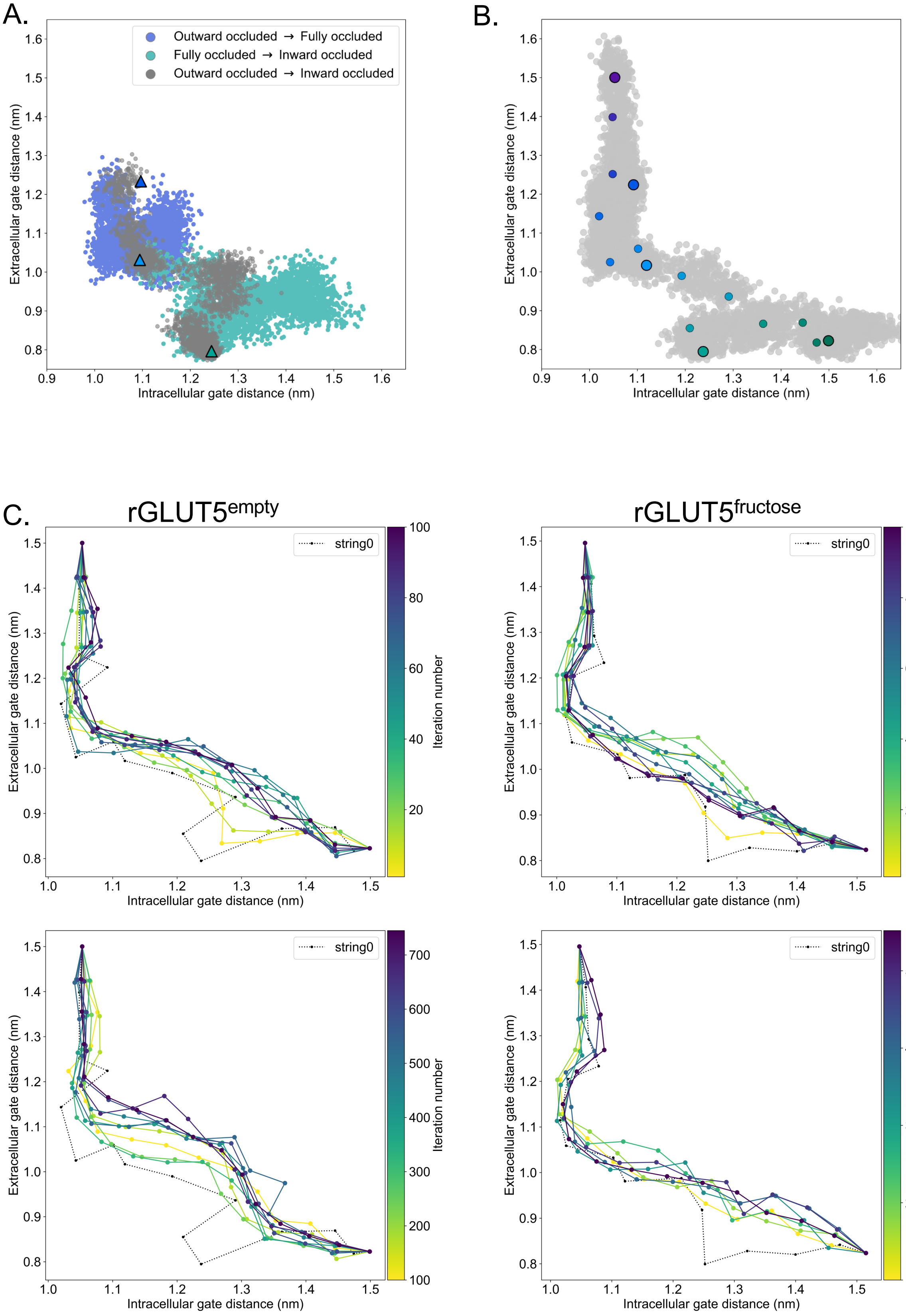
A.
Outward occluded → Fully occluded
Fully occluded → Inward occluded
Outward occluded → Inward occluded
Extracellular gate distance (nm)
Intracellular gate distance (nm)
B.
Extracellular gate distance (nm)
Intracellular gate distance (nm)
C.
rGLUT5empty
rGLUT5fructose
string0
Iteration number
Extracellular gate distance (nm)
Intracellular gate distance (nm)

**Figure S3 – Details of TMD, string simulation setup and convergence. A.** rGLUT5$^{empty}$ TMD with and without the occluded state as a target. Coloring as in rGLUT5$^{fructose}$ TMD results, Fig. 2C. **B.** Beads chosen for the string simulations from the TMD along the CV space for rGLUT5$^{empty}$. The cloud of grey dots represent all CV configurations through the TMD simulations, larger colored dots represent each initial homology model, and the smaller colored dots represent the beads between each of these models, which were chosen for the first iteration of the string-of-swarms method. Beads for rGLUT5$^{fructose}$ found in Fig. 2D. **C.** String convergence as measured by the change of CV values (gate distances) per iteration. Top, before convergence, iterations 1-100, with every 10$^{th}$ iteration shown. Bottom: after convergence, iterations 100-745 (rGLUT5$^{empty}$) or 100-552 (rGLUT5$^{fructose}$), with every 80$^{th}$ iteration shown. The initial string was chosen along the beads highlighted in panel B and Fig. 2D respectively, indicated with a dotted black line labelled ‘string0’.

### Figure S4

A.

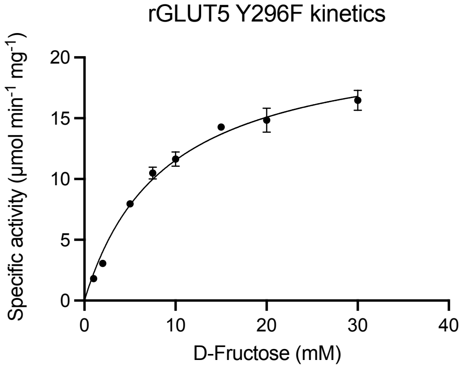


| | WT * | Y296F |
|---|---|---|
| $K_m$ (mM) | 10.91 | 8.8 |
| $V_{max}$ (μmol·min$^{-1}$·mg$^{-1}$) | 52.30 | 21.75 |
| $K_{cat}$(s$^{-1}$) | 43.28 | 48.3 |

B.

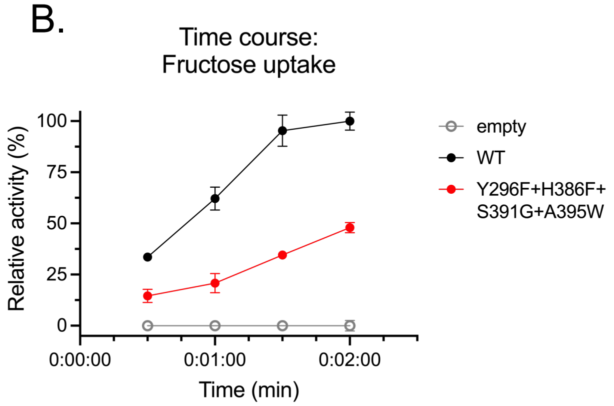


C.

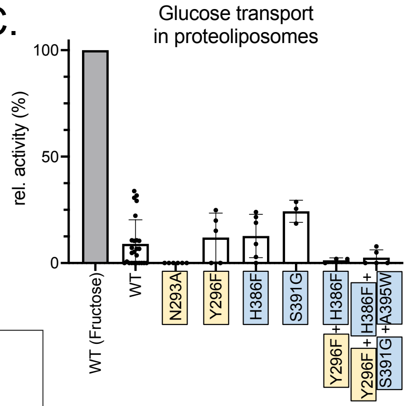


D.

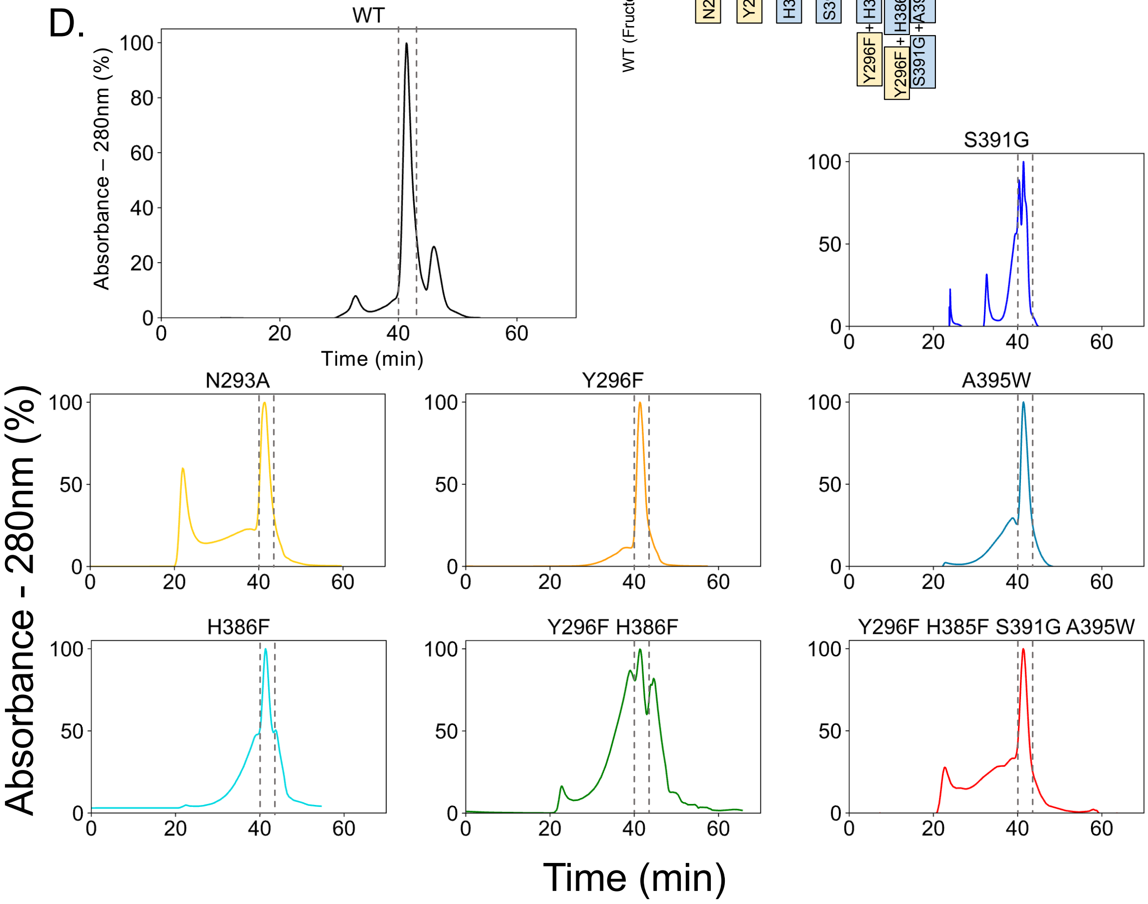

**Figure S4 – GLUT5 kinetics and SEC traces for purified rGLUT5 proteins. A. Zero trans** kinetics (influx) of the rat GLUT5 mutation Y296F. Error bars represent the SEM of n = 3 individual experiments. The $K_M$ is an average calculated from 2 independent reconstitutions and titrations. The kinetic parameters for rat GLUT5 WT (indicated by *) originate from our previously measured kinetics[1]. **B.** Time course of low-activity rGLUT5 mutations (Fig. 4C), normalized between 100% (signal from WT transport after 2 minutes) and 0% (noise from empty liposomes as background). Error bars represent the SEM of n = 3 individual experiments. **C.** $^{14}$C D-glucose transport activity of WT rGLUT5 and mutations as seen in Figure 4B. normalized between 100% (signal from WT fructose transport after 2 minutes) and 0% (noise from empty liposomes as background). Errors bars represent s.e.m. of at least 3 independent measurements taken from at least 1 independent liposome reconstitution. **D.** Relative UV absorbance of DDM-purified rGLUT5 proteins during size exclusion chromatography of the final purified sampled. The rGLUT5 peak between grey dashes was collected for proteoliposome transport assays. Each curve is normalized to the maximum intensity of each trace as 100%.

Figure S5

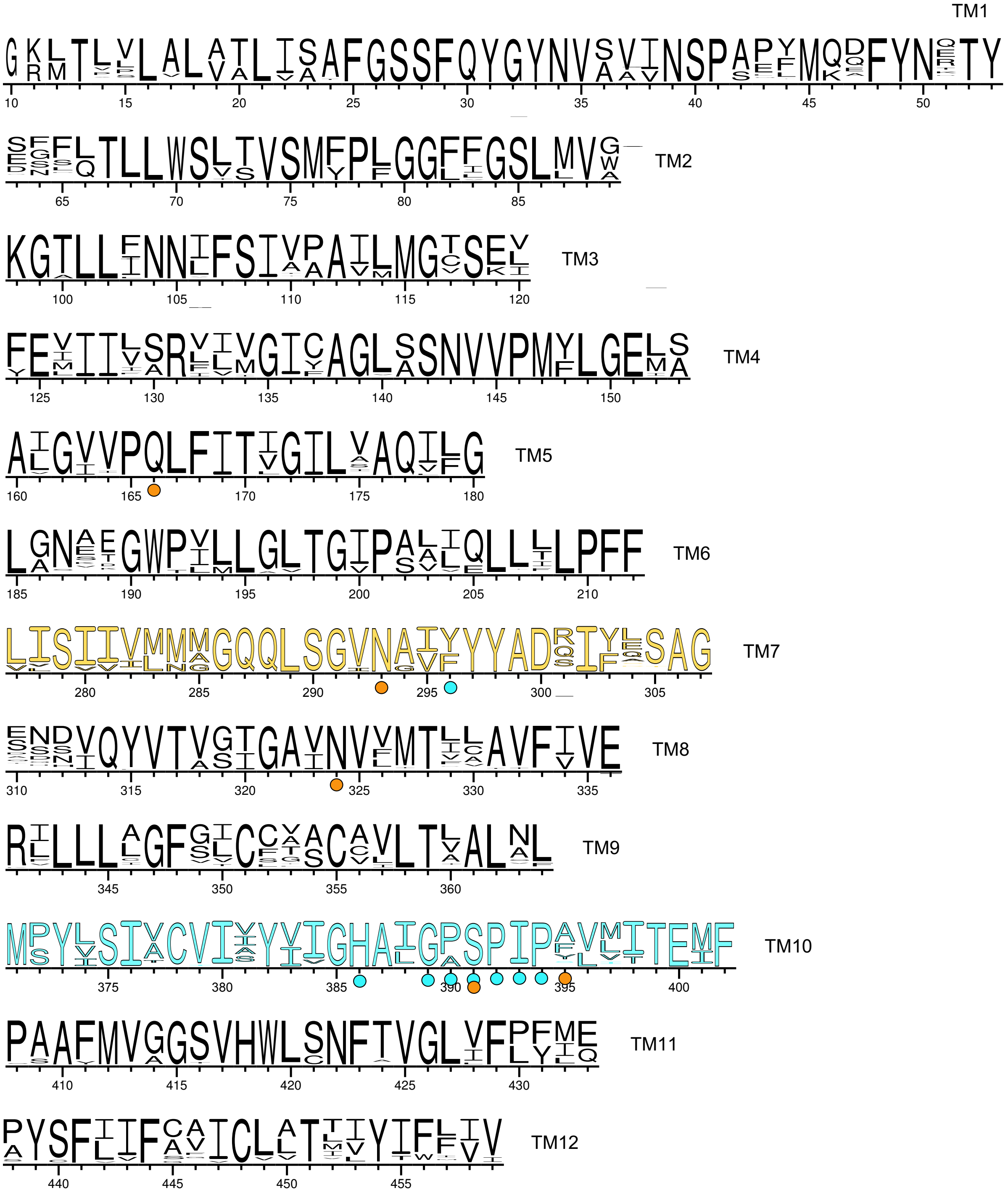

**Figure S5 – Sequence logos of putative GLUT5 proteins.** Sequence logos for transmembrane helices of putative GLUT5 proteins (see Methods) in other organisms. Column numbering corresponds to *rat* GLUT5 residue numbers. TM7 and TM10 helices are colored as in Figure 1. Following coloring as in Figure S1, residues involved in fructose binding are indicated with an orange circle, the GPXPXP motif as well as residues involved in coupling fructose binding to conformational change are indicated with a cyan circle. The TM7b tyrosine or phenylalanine can be found at column 296.

### Figure S6

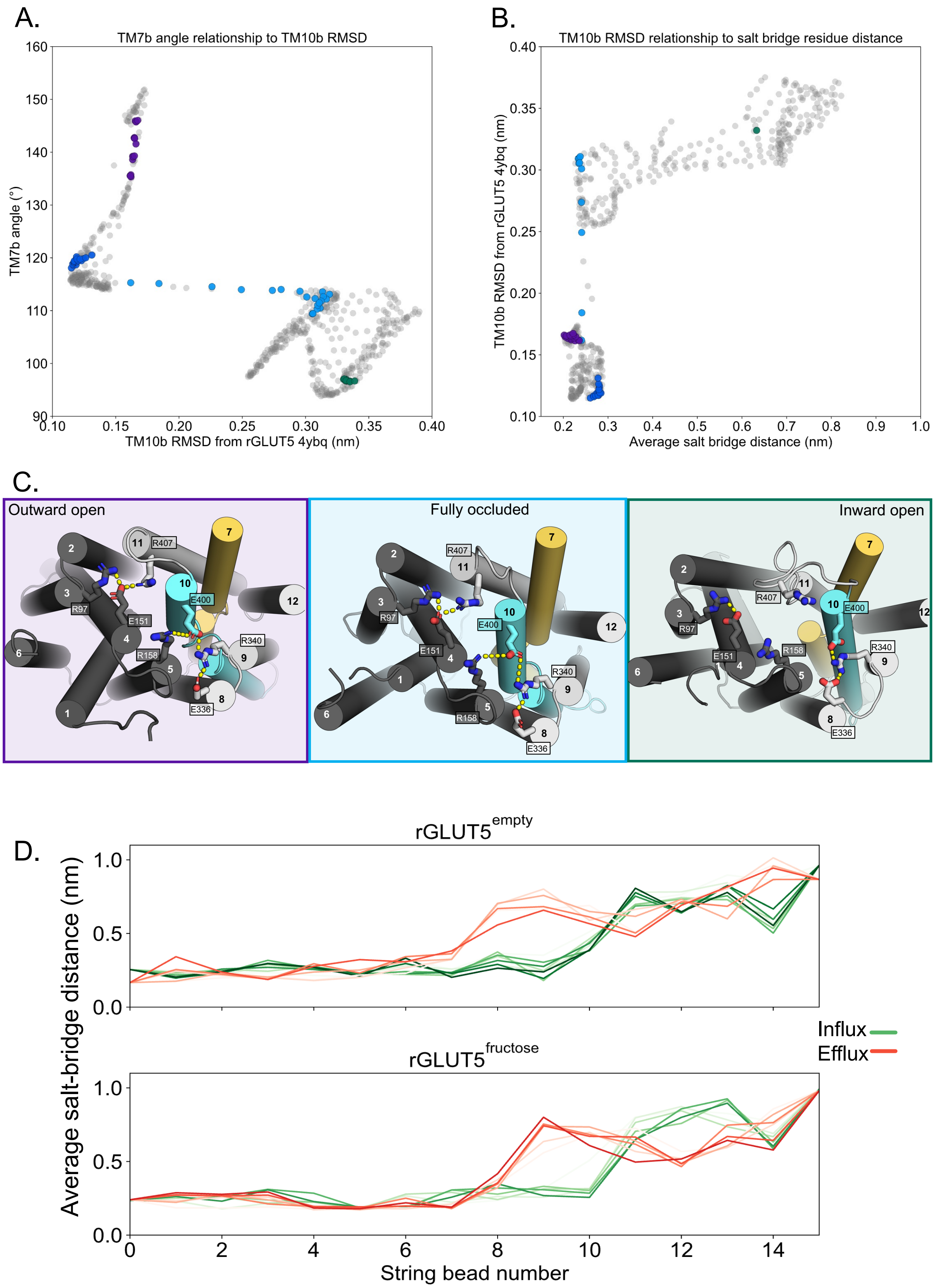
A.
TM7b angle relationship to TM10b RMSD
TM7b angle (°)
TM10b RMSD from rGLUT5 4ybq (nm)
B.
TM10b RMSD relationship to salt bridge residue distance
TM10b RMSD from rGLUT5 4ybq (nm)
Average salt bridge distance (nm)
C.
Outward open
Fully occluded
Inward open
R407
E400
R97
E151
R158
R340
E336
D.
rGLUT5^empty
rGLUT5^fructose
Average salt-bridge distance (nm)
String bead number
Influx
Efflux

**Figure S6 – Salt bridge formation during the conformational cycle. A.** Relationship between TM7b angle and TM10b RMSD. Circles colored correspond to bins with EC and IC values near the outward open (purple), outward occluded (deep blue), occluded (light blue), and inward open (green) conformations, respectively. The trajectory does not sample enough around the original inward occluded model to enable its inclusion in the analysis. **B.** Relationship between TM10b RMSD and the average of the two state-dependent salt bridge network distances. Coloring as in panel A. **C**. The salt bridge residues of GLUT5 in three of the major conformations, forming two main salt bridge groups, where interactions are indicated by yellow dashes. State-dependent interactions are observed for the salt bridge between E151-R401 (TM4-TM11) and E400-R158 (TM10-TM5), which are not observed in the inward open state (right, green). Protein coloring as in Figure 1, and background coloring indicates states representing the same colors in panel A and B. **D.** Measuring the average of the two state-dependent salt bridge distances during the transport cycle (panel C) in several beads, through iterations. Earlier iterations are shown in a light shade, and the final iteration is darkest (rGLUT5$^{empty/\ influx}$: 745, rGLUT5$^{empty/\ efflux}$: 450, rGLUT5$^{fructose/\ influx}$: 552, rGLUT5$^{fructose/\ efflux}$: 553), with every 50$^{th}$ iteration in between displayed.

### Figure S7

A.

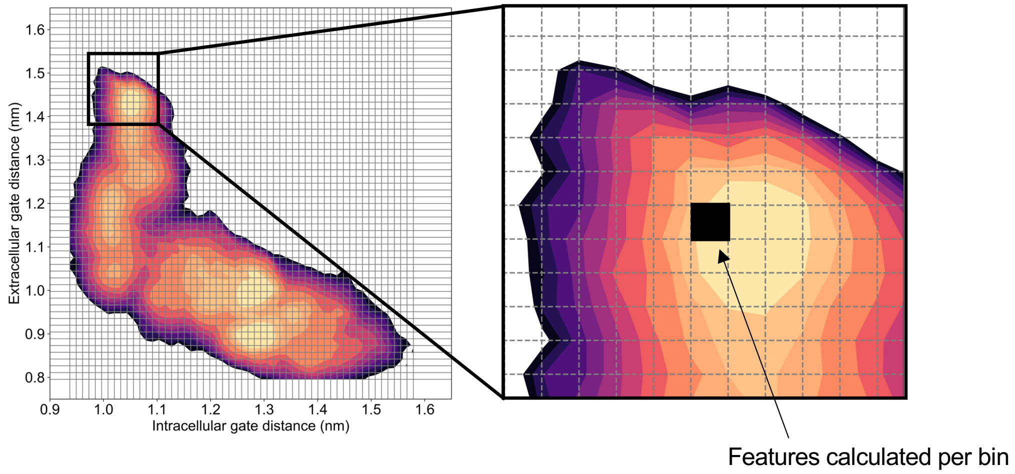


B.

rGLUT5$^{empty}$

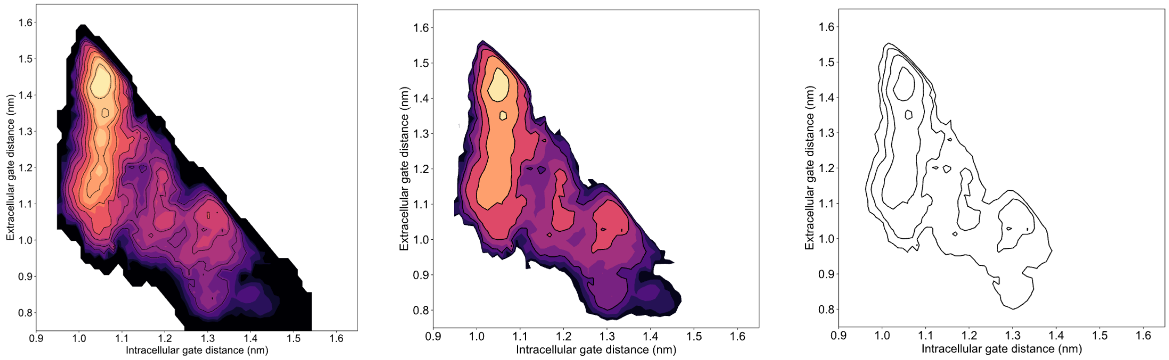


rGLUT5$^{fructose}$

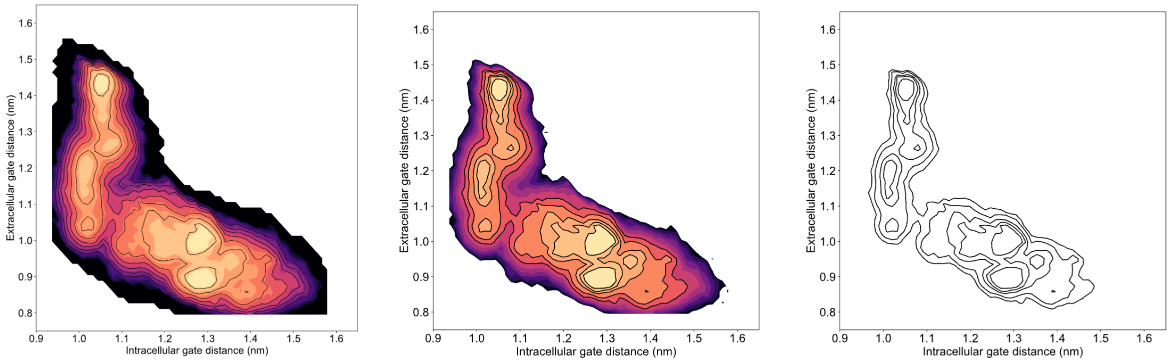

**Figure S7 – Calculation of features along energy surfaces**. **A.** A simplified graphic of how each bin is extracted from the free energy surface. **B.** A simplified graphic of how the black line abstractions used for analysis originate from the free energy surfaces for Fig. 3A, Fig. 4C.

1 Qureshi, A. A. *et al.* The molecular basis for sugar import in malaria parasites. *Nature* **578**, 321-325 (2020). https://doi.org:10.1038/s41586-020-1963-z
